# Modeling of Aberrant Epithelial Reprogramming in Idiopathic Pulmonary Fibrosis using Human Induced Pluripotent Stem Cell-derived Alveolar Organoids

**DOI:** 10.1101/2022.06.15.496102

**Authors:** Victoria Ptasinski, Susan J. Monkley, Karolina Öst, Markus Tammia, Catherine Overed-Sayer, Petra Hazon, Darcy E. Wagner, Lynne A. Murray

## Abstract

Repeated injury of the lung epithelium is proposed to be a main driver of idiopathic pulmonary fibrosis (IPF). However, none of the available therapies target the epithelium and there is a limited amount of human models of fibrotic epithelial damage with suitability for drug screening and discovery. We developed a model of the epithelial reprogramming seen in IPF using alveolar organoids derived from human induced pluripotent stem cells stimulated with a cocktail of pro-fibrotic and inflammatory cytokines. This fibrosis cocktail induced persistent epithelial reprogramming and expression of extracellular matrix. Deconvolution of RNA-seq data indicated that the fibrosis cocktail increased the proportion of cells with the *KRT5^-^/KRT17^+^* aberrant basaloid phenotype, recently identified in the lungs of IPF patients. Treatment with nintedanib and pirfenidone had effects on markers of extracellular matrix, pro-fibrotic mediators and epithelial reprogramming. Thus, our system recapitulates key aspects of IPF and is a promising system for drug discovery.

## Introduction

Idiopathic pulmonary fibrosis (IPF) is a chronic lung disease with a median survival of 2-3 years (1) characterized by thickening of the alveolar walls and extracellular matrix (ECM) deposition in the distal part of the lung (2). The available therapies, nintedanib and pirfenidone, slow down disease progression but are not curative (3). Current knowledge suggests that repeated alveolar epithelial injury is a driver of disease (4). Functional repair is impeded in IPF patients leading to loss of alveolar type 1 (AT1) and type 2 (AT2) cells. The lung epithelium in patients with IPF is increasingly recognized as dysregulated and cell populations with transient or indeterminate phenotypes appear (5) due to aberrant epithelial reprogramming. This includes the appearance of airway epithelial-like cells in the alveoli (6) and the epithelial-to-mesenchymal transition (EMT) state where epithelial cells acquire mesenchymal markers (7). Recent studies have identified subpopulations of epithelial cells with increased pro-fibrotic transcriptional activity in the lungs of IPF patients (8–12). This includes the aberrant basaloid cells, which co-express markers of airway basal cells, mesenchyme, and senescence (9, 10), and the transient AT2-AT1 state (11, 12). However, the functional contribution of these aberrant subpopulations to the IPF pathogenesis has not been studied since it is not presently known what induces this phenotypic change. Thus, there is a lack of models that mimic this aspect of epithelial reprogramming in IPF. Despite the urgent need for such models, the access to relevant pre-clinical systems in which expandable human AT2 cells are utilized is limited. Although organoid models have enabled maintenance of the AT2 phenotype in culture (13), the poor proliferation of primary AT2 cells after isolation from the lung remains an issue. To address this obstacle, several groups have reported advances in deriving proliferative and functional AT2 cells from human embryonic stem cells (ESCs) (14) or induced pluripotent stem cells (iPSCs) (15–18). These cells have been used in modeling genetic lung diseases such as Hermansky-Pudlak Syndrome-associated interstitial pneumonia (HPSIP) (19, 20), complex chronic lung diseases like pulmonary fibrosis (21–24) and responses to environmental injurious stimuli (25). However, there is still a lack of models of fibrotic epithelial reprogramming which are adaptable for drug screening. Such models should include the use of expandable cells which can be stored as frozen banks to allow for high throughput drug screening and cultured in a miniaturized format with stimuli inducing persistent disease-relevant changes.

In this study, we developed a model of fibrotic alveolar epithelial reprogramming suitable for drug discovery. iPSC-derived AT2 (iAT2) cells from frozen banks were exposed to a cocktail of pro-fibrotic and inflammatory cytokines which has been shown to induce early fibrotic changes and alterations in epithelial cell phenotypes in human precision-cut lung slices (PCLS) and has been useful for evaluation of anti-fibrotic compounds in the same system (26, 27). We describe that stimulation of iAT2 cells with this fibrosis cocktail induces persistent reprogramming into a similar aberrant basaloid-like state as reported in IPF and production of ECM. We also show that our model is responsive to treatment with nintedanib and pirfenidone, demonstrating that this system can be adapted for drug screening. Thus, this system models several aspects of human IPF allowing for studies of fibrosis induction and meets the desirable criteria for application in drug discovery.

## Results

### Fibrosis cocktail induces transcriptomic changes linked to IPF

To create an expandable and biologically relevant cell source for modeling alveolar epithelial injury in IPF, we differentiated AT2 cells from iPSCs using a method based on previously described protocols (17, 18, 28) (Fig. S1A). As the published protocols utilized other iPSC lines, we first validated the efficiency of these protocols in our cell line. Gating for selection of CD47^high^/CD26^low^ cells enriched for putative lung progenitors expressing NKX2.1, as previously reported (Fig. S1B-C) (28). A 3-day definitive endoderm (DE) induction combined with cell subculturing at day 8 of differentiation provided the highest yield of CD47^high^/CD26^low^ cells with the iPSC line used in our study (Fig. S1D). Subculturing during lung progenitor induction has been suggested to improve the yield in certain cell lines (17). Since we could not see a significant difference between splitting the cells at a ratio of 1:3 and 1:6, we continued with the 1:3 ratio to avoid the sparseness of the cells caused by the 1:6 ratio split. Organoids of iAT2 cells (alveolospheres) expressed the surfactant protein SP-B in intracellular granules likely associated with lamellar bodies and as a secreted form (Fig. S1E) which is a feature specific to AT2 cells (29), thus confirming that our cell line and culture conditions enabled us to generate iAT2 cells using the previously described protocols.

As our goal was to build a model suitable for drug screening, we next evaluated the ability to cryopreserve alveolospheres to allow for timing control of the screens. Therefore, we assessed the AT2 phenotype before and after cryopreservation using standard cell freezing conditions containing DMSO to determine if the alveolospheres can be stored as frozen banks. We found that the expression of another AT2-cell specific marker pro-surfactant protein C (pro-SP-C) (30) was retained for at least 2 passages after cryopreservation, thawing and expansion, at which point the cells reached a suitable number for experiments (Fig. S1F).

Next, we explored the possibility to induce responses linked to IPF in the alveolospheres. Alveolospheres, formed from single iAT2 cells over 14 days, were stimulated for 72 h with a fibrosis cocktail (FC) containing transforming growth factor β (TGF-β), tumor necrosis factor α (TNF-α), platelet-derived growth factor AB (PDGF-AB) and lysophosphatidic acid (LPA), previously shown to induce phenotypic changes of epithelial cells in an *ex vivo* model mimicking alterations seen in IPF (26, 27) (Fig. 1A). We found that the majority of alveolospheres exhibited a ‘dense’ morphology after FC stimulation compared to the control cocktail (CC) containing the diluents, where the alveolospheres retained a visible lumen (Fig. 1B). To gain insight into the transcriptomic changes induced by the FC, we performed RNA-seq of alveolospheres stimulated with the FC or CC. Principal component analysis showed clear separation of samples based on the stimulation, explaining 84.7 % of the variance (Fig 1C). The top 50 differentially expressed genes by fold change (Fig. S2A) included downregulation of the AT2-cell marker *SFTPC* (30) and upregulation of the fibrosis-associated genes *MMP7* (31) and *SERPINE1* (32) related to ECM remodeling and senescence. For all differentially expressed genes, see List S1. The genes induced by the FC were associated with biological processes linked to IPF such as ECM organization (33), cell adhesion (34), and inflammatory or immune responses (35) (Fig. 1D). This set of genes was associated with extracellular and cell membrane-associated structures, such as focal adhesions which are involved in regulation of myofibroblast survival upon stimulation with TGF-β1 (36) (Fig. 1E). The upregulated genes are also known to engage in cell-cell interactions such as binding of cadherin, integrin and actin (Fig. 1F). Pathway annotation of the upregulated genes using the Kyoto Encyclopedia of Genes and Genomes (KEGG) demonstrated association with several pathways previously reported to be related to IPF (Fig. 1G) including the NF-κβ and MAPK signaling pathways (37, 38), the Hippo/YAP signaling pathway which is altered in epithelial cells in IPF (39), and the p53 signaling pathway related to apoptosis and senescence (40). Epithelial cells have been previously shown to express factors of the senescence-associated secretory phenotype (SASP) in experimental pulmonary fibrosis and in IPF tissue (41, 42), several of which were also affected by the FC in our model (Fig. S2B). On the other hand, genes downregulated by the FC were associated with processes such as DNA replication, cell division, mitotic nuclear division, and cell cycle transition (Fig. 1D). We also found associations with telomere maintenance and DNA repair which are known to be impaired in IPF (43) (Figure 1D and Fig. S2C). In contrast to the upregulated genes, the downregulated genes associated more with intracellular structures such as ribosomes, nucleus and lamellar bodies, important for surfactant processing in AT2 cells (29) (Fig. 1E) and with functions and pathways particularly linked to ribosomes, DNA replication and metabolism (Fig. 1F and G). For all gene ontology results, see Lists S2-S9.

**Figure 1:**
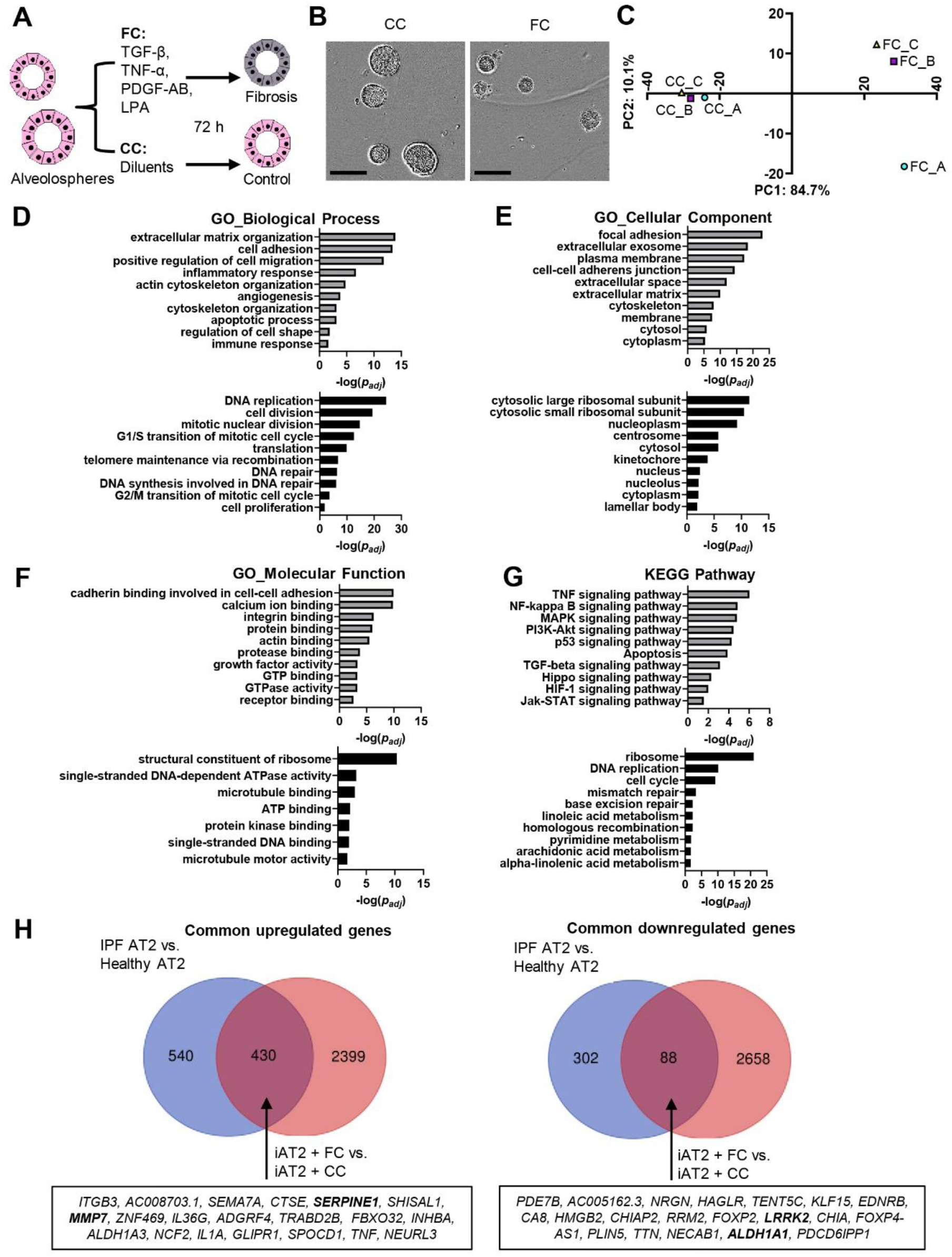
Fibrosis cocktail induces transcriptomic changes linked to IPF. A) Schematic showing FC stimulation of alveolospheres. B) Representative phase contrast images of alveolospheres stimulated with the CC or FC for 72 h. 4x magnification. Scale bar = 250 µm. C) PCA plot of RNA-seq data, top 1000 transcripts by coefficient of variation. *n* = 3 batches of iAT2 cells. D-G) Gene enrichment analyses of the upregulated (grey) and downregulated (black) genes according to their gene ontology (GO) biological process, GO cellular component, GO molecular function and associated Kyoto Encyclopedia of Genes and Genomes (KEGG) pathway. H) Venn-diagrams of common dysregulated transcripts in FC-stimulated alveolospheres and primary AT2 cells isolated from IPF patients (GEO: GSE94555). Selected top 20 dysregulated genes (by absolute fold change) are denoted in the boxes. For D-H: genes selected by *p(adj)* < 0.05, absolute log2 fold change ≥ 0.7. See also Fig. S1 and S2 and Lists S1-S11.

Next, we sought to compare the similarities in the genes altered by the FC with those seen in IPF patients. To do this we compared our data with a public dataset of primary AT2 cells derived from IPF patients and normal donors (5). Several fibrosis-associated markers were upregulated in both primary fibrotic AT2 cells and the FC-stimulated alveolospheres, including *MMP7* and *SERPINE1* (Fig 1H and List S10). Among the common downregulated transcripts (Fig. 1H and List S11), we found *LRRK2* which loss is linked to AT2 cell dysfunction in experimental pulmonary fibrosis (44). We also noted *ALDH1A1* in this group of genes, which is associated with stemness and exert anti-fibrotic properties when highly expressed in lung epithelial cells in experimental pulmonary fibrosis models in mouse (45). Taken together, these results show that the FC-induced transcriptomic changes share similarities to those seen in IPF.

### Fibrosis cocktail induces epithelial injury

As the loss of alveolar cell phenotypes is a prominent characteristic of IPF, we sought to determine whether the FC induces such responses in the alveolospheres. Consistent with previously published observations in human PCLS (26), we observed downregulation of the AT2 cell markers *SFTPC* and *SFTPB* upon FC stimulation (Fig. 2A-C). The decrease of the processed, mature form of SP-B at around 6 kDa (Fig. 2B) indicates loss of the functional AT2 phenotype upon FC stimulation (29). We were curious to see if the loss of the AT2 phenotype was associated with a similar EMT state as seen in IPF (7). We observed increased mesenchymal *CDH2* expression (Fig. 2A) and VIM expression localized to individual cells (Fig. 2C), although this was not associated with a loss of the epithelial marker *CDH1*/ECAD indicating that the cells did not undergo a full EMT transition (Fig. 2A, C). This data suggests that the stimulation with the FC induces AT2 cell dysfunction and an aberrant epithelial cell phenotype.

**Figure 2:**
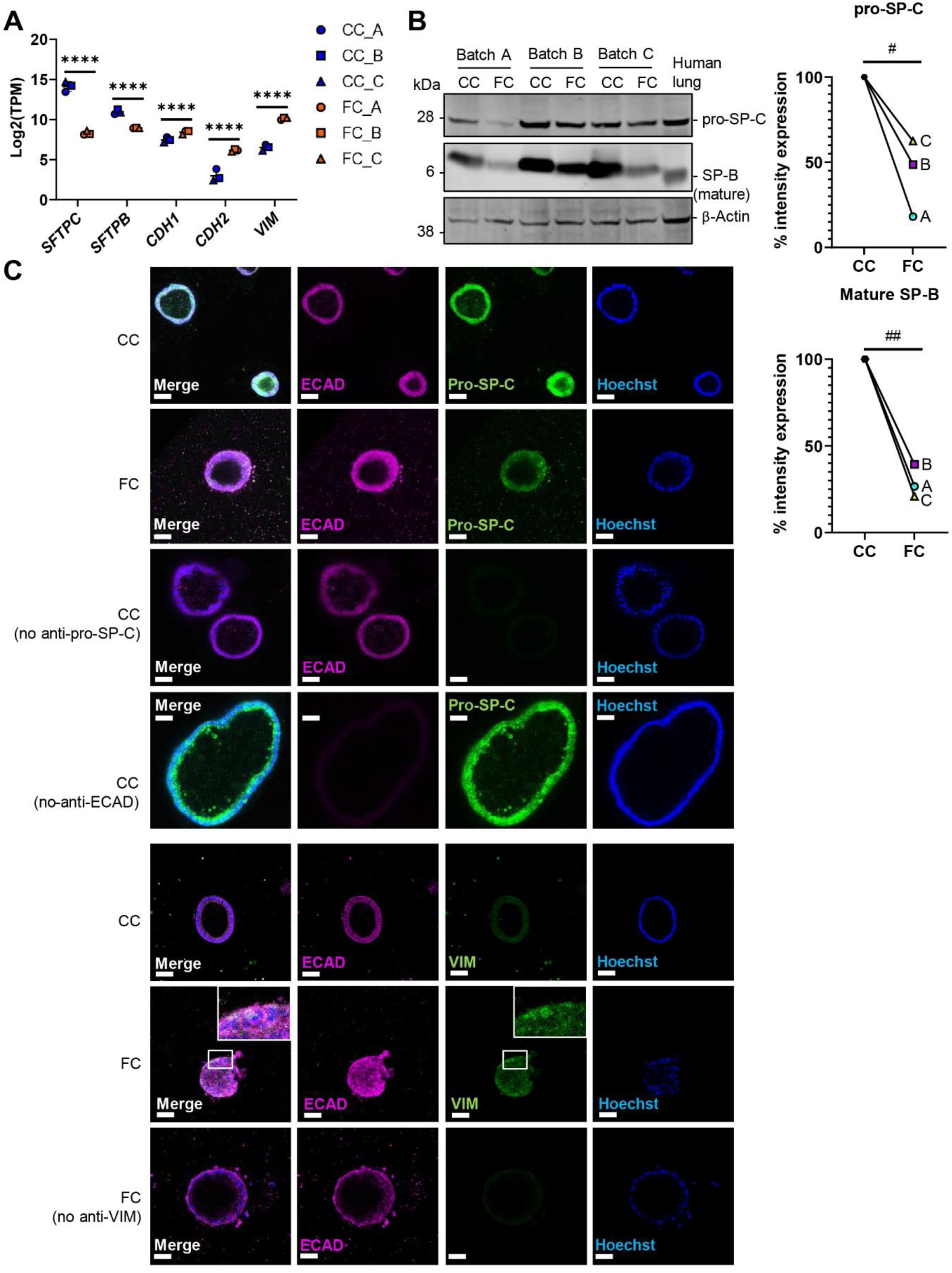
Fibrosis cocktail induces epithelial injury. A) Expression of selected epithelial and reprogramming-associated genes. Data presented as means ± SD. **** = *p(adj)* < 0.0001. *n* = 3 batches of iAT2 cells. B) Western blot of intracellular protein lysates from alveolospheres stimulated with CC or FC. Human lung lysate as positive control. Signal was quantified as band intensity normalized to β-actin and expressed as percentages compared to the respective CC sample set to 100 % expression. ## = *p* < 0.01, # = *p* < 0.05 by one sample t-test on the percentage intensity compared to a hypothetical value of 100. *n* = 3 batches of iAT2 cells. Blots are cropped from the same membrane and are obtained by sequential blotting as outlined in Supplemental Experimental Procedures. Brightness and contrast has been enhanced. C) Immunofluorescence of alveolospheres stimulated with CC or FC. No anti pro-SP-C, anti-ECAD and anti-VIM as negative controls. Brightness and contrast has been enhanced. Scale bar = 50 µm.

### Fibrosis cocktail induces epithelial reprogramming into an aberrant basaloid phenotype

After observing signatures of epithelial reprogramming induced by the FC stimulation based on expression of individual markers, we were curious to explore changes in phenotypes associated with cell types. For this, we used computational deconvolution with Bisque (46, 47). For the analysis, we used a public single-cell RNA-seq dataset (GEO Accession: GSE135893) as a reference filtered to include lung epithelial cells from non-fibrotic controls and IPF donors (10) (Fig. 3A). Deconvolution of our alveolosphere RNA-seq dataset with Bisque estimated a decreased proportion of cells with the AT2 phenotype by the FC stimulation (Fig. 3B, Fig. S3A), which is consistent with our earlier observations. Intriguingly, the deconvolution estimated an increased proportion of cells with the *KRT5^-^/KRT17^+^* aberrant basaloid phenotype by the FC stimulation (Fig. 3B). In line with this, we confirmed the upregulation of *KRT17* and downregulation of *KRT5* induced by the FC stimulation in our dataset (Fig. 3C), and detected the appearance of KRT17^+^ cells upon FC stimulation which was most prominent in the organoids with a dense morphology (Fig. 3D). We also noted FC-induced expression of *KRT8* characterizing the transient AT2-AT1 cell population in IPF (11) (Fig. 3C), which shares transcriptional similarity with the *KRT5^-^/KRT17^+^*aberrant basaloid cells (12). In agreement with previously published observations, the KRT8 expression was localized to the KRT17^+^ cells (12, 48) upon stimulation with the FC (Fig. 3D). The deconvolution also predicted changes in other cell type proportions such as ciliated and *SCGB3A2^+^* cells (Fig. S3A) although these were contradictory to the expression patterns of the classical cell type markers in our dataset (Fig. S3B). This is likely due to the fact that Bisque uses a signature of genes rather than individual markers to determine the cell types (46). Altogether, our data shows that the FC induces reprogramming in a subpopulation of cells in our model into an aberrant basaloid-like phenotype, similar to what is seen in lungs of IPF patients (9, 10).

**Figure 3:**
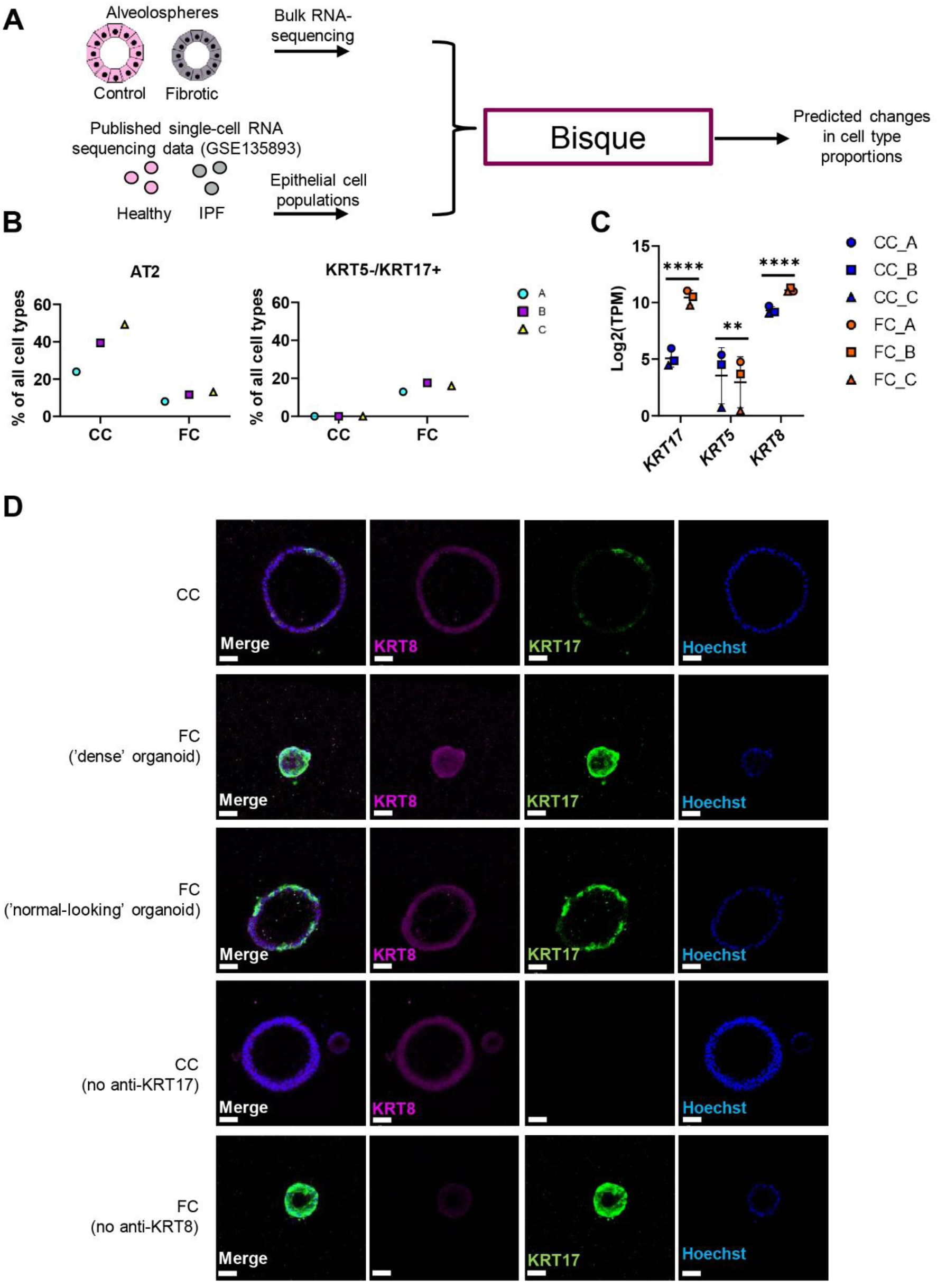
Deconvolution analysis predicts increase in aberrant basaloid like-cells by the fibrosis cocktail. A) Schematic depicting Bisque reference-based analysis of epithelial cell populations from non-fibrotic controls and IPF from the publicly available single-cell RNA dataset (GEO Accession: GSE135893). B) Selected epithelial cell proportions estimated by Bisque in alveolospheres stimulated with CC or FC. *n* = 3 batches of iAT2 cells. C) Expression of selected epithelial genes in alveolospheres. Data presented as means ± SD. **** = *p(adj)* < 0.0001, ** = *p(adj)* < 0.01. *n* = 3 batches of iAT2 cells. D) Immunofluorescence of alveolospheres stimulated with CC or FC. No anti-KRT8 and anti-KRT17 as negative controls. Brightness and contrast has been enhanced. Scale bar = 50 µm. See also Fig. S3.

### Fibrosis cocktail induces production of ECM

The single-cell studies describing the *KRT5^-^/KRT17^+^* aberrant basaloid cells have shown that these cells express markers of ECM (9, 10). We therefore assessed the transcriptional changes of matrisome components induced by the FC (Fig. 4A) (49). Expression of several matrisome genes was altered by the FC, including upregulation of the genes *FN1*, *COL1A1* and *TNC* expressed by the aberrant basaloid cells (Fig. 4B) (9, 10). We also observed increased secretion of the ECM proteins fibronectin (FN), tenascin C (TNC) and pro-collagen 1a1 (COL1α1) induced by the FC in the medium (Fig. 4C-E). As collagen is a major component of the deposited ECM in IPF, we examined the localization of the collagen 1 (COL1) expression in alveolospheres (Fig. 4F). We observed that the deposition of COL1 in the FC-stimulated alveolospheres was mostly visible on the periphery of the organoids, indicating that the cells have a polarized secretion of COL1 on the basal side. In line with our earlier observations, this was not associated with loss of the epithelial marker ECAD. This data shows that the FC stimulation induces production of ECM from epithelial cells.

**Figure 4:**
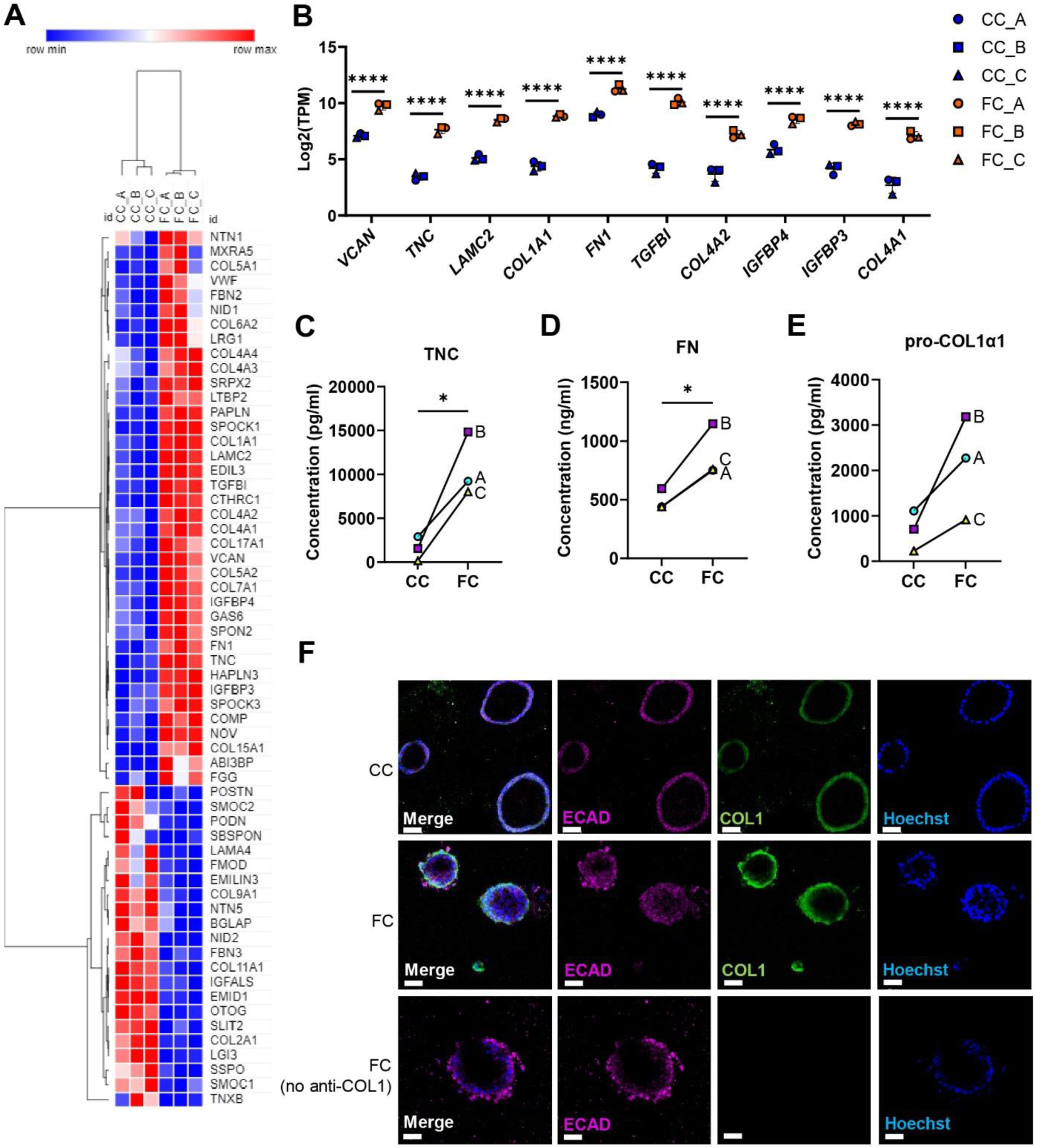
Fibrosis cocktail induces production of extracellular matrix. A) Heatmap of significantly dysregulated matrisome genes in FC-stimulated alveolospheres by absolute fold change (*p(adj)* < 0.05). B) Selected matrisome genes with the highest expression in FC-stimulated alveolospheres (*p(adj)* < 0.05, log2(TPM) ≥ 5, log2 fold change ≥ 2). Data presented as means ± SD. **** = *p(adj)* < 0.0001. *n* = 3 batches of iAT2 cells. C) Secreted concentrations of tenascin C measured by ELISA. * = *p* < 0.05 by paired, two-tailed t-test. D) Secreted concentrations of fibronectin measured by ELISA. * = *p* < 0.05 by paired, two-tailed t-test. E) Secreted concentrations of pro-collagen 1α1 measured by ELISA. Paired, two-tailed t-test performed. F) Immunofluorescence of alveolospheres stimulated with CC or FC. No anti-COL1 included as negative control. Brightness and contrast has been enhanced. Scale bar = 50 µm.

### Fibrotic responses persist after withdrawal of the fibrosis cocktail

A desired feature of an experimental model of IPF is sustainment of the fibrotic responses over time. Thus, we assessed if the changes we observed in our initial experiments related to ECM production, senescence and aberrant epithelial reprogramming seen after 72 h (3 days) of FC stimulation (Fig. S4A) persisted after withdrawal of the FC. After the stimulation for 3 days, we replaced the FC or CC medium with the normal CK+DCI maintenance medium (Table S1 and S4) (17, 18) for 4 additional days (Fig. 5A) to see if the changes we induced via FC persisted or returned to the levels seen in their time matched CC control (Fig. S4B). A majority of the FC-stimulated alveolospheres still exhibited a dense morphology after the withdrawal (Fig. 5B). Thus, we continued with exploring the gene expression of selected markers for which we observed significant or near significant (*p* < 0.07) changes at day 3 of FC stimulation (Fig. S4A). We first assessed if the ECM production was sustained after the FC withdrawal, and found that withdrawal of the FC did not restore the gene expression or protein secretion in the medium of any of the ECM markers *FN1, TNC* and *COL1A1* to their time matched CC controls (Fig. 5C-D). Next, we assessed whether the alveolospheres showed signs of alveolar recovery after the FC withdrawal. We assessed the genes associated with senescence (*CDKN1A* and *CDKN2A*) and found no restoration of the expression to their time matched CC controls following the FC withdrawal (Fig. 5E), indicating persistent senescence of the cells. We further explored the expression of genes associated with the alveolar epithelial phenotype and reprogramming (Fig. 5F). Although we could not see a significant increase of the surfactant protein *SFTPC* following the FC withdrawal, we did observe a significant increase in the expression of *SFTPB* (Fig. 5F) indicating attempt of repair by the cells. Intriguingly, the aberrant basaloid cell-associated markers *KRT8* and *KRT17* (Fig. 5F-G) (9, 10, 12) were still elevated in the FC-stimulated alveolospheres after withdrawal, suggesting that this aberrant reprogramming was non-reversible within the assessed time frame. This data shows that stimulation with the FC induces sustained pro-fibrotic responses, which allows longer studies of fibrotic epithelial injury and reprogramming.

**Figure 5:**
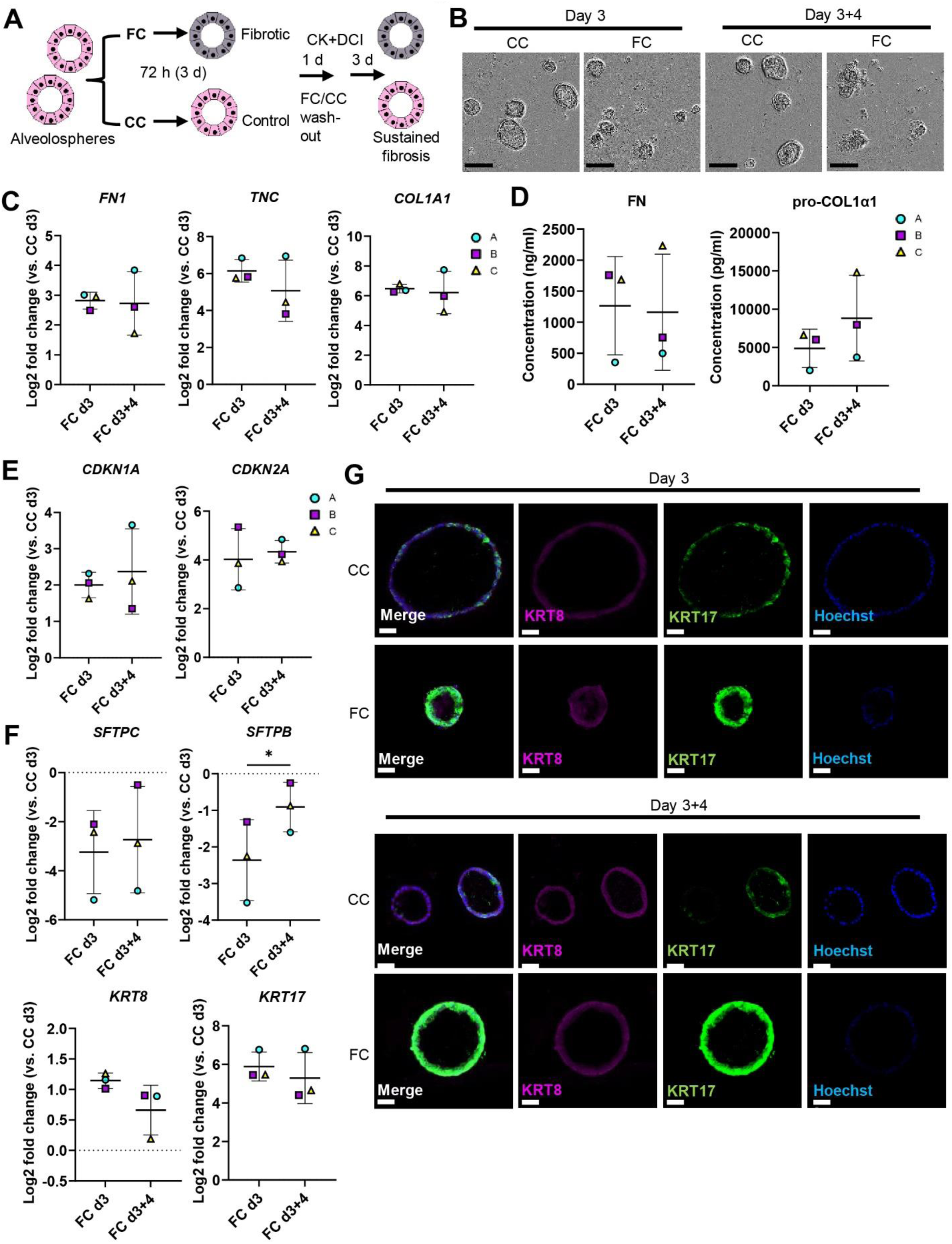
Effects of fibrosis cocktail persist after withdrawal. A) Schematic showing FC stimulation of alveolospheres to assess persistence of fibrotic responses. B) Representative phase contrast images of alveolospheres stimulated with the CC or FC for 3 days and after withdrawal of the CC or FC for 4 days. 4x magnification. Scale bar = 250 µm. C) Expression of genes measured by qRT-PCR related to extracellular matrix production. D) Secreted protein concentrations of fibronectin and pro-collagen 1α1 in the medium measured by ELISA. Data presented as means ± SD. Paired, two-tailed t-test performed. *n* = 3 batches of iAT2 cells. E) Expression of genes measured by qRT-PCR related to senescence. F) Expression of genes measured by qRT-PCR related to alveolar epithelial injury and reprogramming. G) Immunofluorescence of alveolospheres stimulated with CC or FC after 3 days of stimulation and following 4 days of withdrawal. Scale bar = 50 µm. Brightness and contrast has been enhanced. For C, E and F: Expression of genes normalized to the average CT value of the reference genes *GAPDH, TBP* and *HPRT1.* Fold changes (2^-ΔΔCT) calculated by comparison to the average ΔCT value of the CC day 3 population. Data presented as means ± SD. Significance tested by paired, two-tailed t-test. *n* = 3 batches of iAT2 cells. See also Fig. S4.

### FC stimulation of iAT2 cells is an adaptable system for drug screening

Next, we sought to assess if our model has potential for adaptation in drug screening. We were interested in testing a treatment approach in which we could potentially preserve the normal phenotype of the AT2 cells in a pro-fibrotic environment, mimicked by the FC. We therefore performed prophylactic treatment of alveolospheres with the anti-fibrotic drugs nintedanib and pirfenidone in combination with the FC stimulation. First, we performed a dose-response study to assay for compound-mediated LDH cytotoxicity (Fig. S5A). Cytotoxicity was not significantly induced after addition of the compounds in concentrations up to 10 µM, although a trend of increase was seen after addition of 10 µM pirfenidone to the FC (Fig. S5B). Thus, higher concentrations of the compounds were not used.

We initially sought to determine which markers were suitable for evaluation of the treatments by looking into the significant changes induced by the FC with the compound vehicle (DMSO) at day 3 (Fig. S6). In line with our earlier observations, we saw significant effects of the FC on markers associated with ECM production (*FN1* and *TNC*) (Fig. S6A-B), pro-fibrotic signaling (*TGFB1*, *MMP7* and *IGFBP3*), senescence (*CDKN2A*) and epithelial reprogramming (*SFTPC*, *SFTPB*, *KRT17, VIM* and *CDH2*) at day 3 (Fig. S6A). Since other *in vitro* studies have reported effects on the expression of the ECM marker *FN1* and the surfactant protein *SFTPC* in murine alveolar epithelial cells grown in 2D tissue culture plastic using nintedanib at 1 μM (27), we initially assessed the responses to concentrations of 0.1 and 1 μM on these markers in our system (Fig. S7A). Unlike these previously published observations, there was no significant effect of the compounds on either of these markers, although there seemed to be a dose-dependent trend of decreased *FN1* expression. We therefore focused on evaluating the effects of the compounds at concentrations of 10 μM (Fig. 6A).

**Figure 6:**
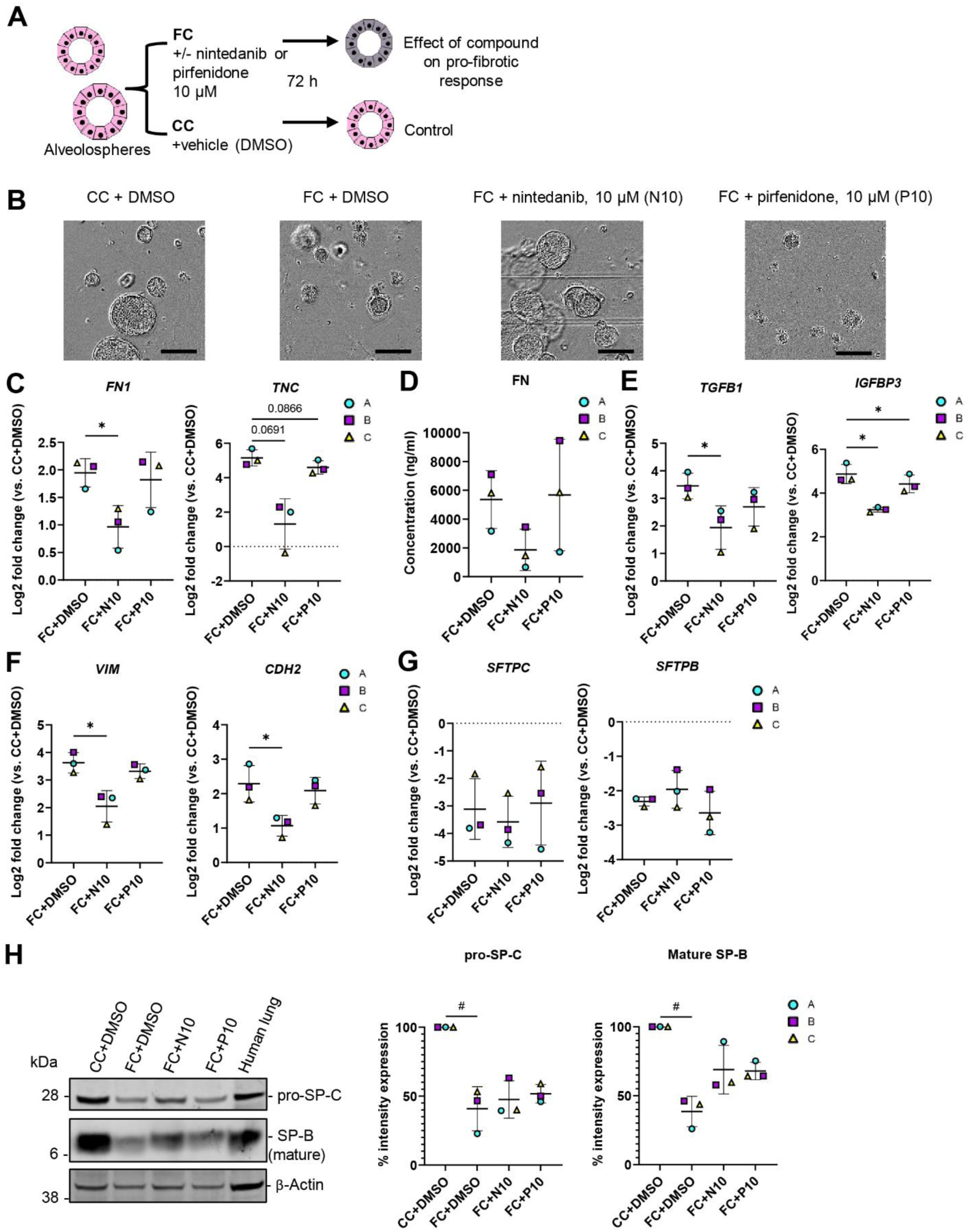
Anti-fibrotic treatment of alveolospheres stimulated with the fibrosis cocktail. A) Schematic depicting prophylactic treatment of FC-stimulated alveolospheres with nintedanib or pirfenidone. B) Representative phase contrast images of alveolospheres stimulated with the CC or FC for 72 h with or without 10 µM nintedanib or pirfenidone. 4x magnification. Scale bar = 250 µm. C) Expression of genes measured by qRT-PCR related to extracellular matrix production. * = *p* < 0.05. D) Secreted protein concentrations of fibronectin in the medium measured by ELISA. Data presented as means ± SD. Repeated measures one-way ANOVA (comparisons to FC+DMSO) performed. *n* = 3 batches of iAT2 cells. E) Expression of genes measured by qRT-PCR related to pro-fibrotic signaling. * = *p* < 0.05. F) Expression of genes measured by qRT-PCR related to alveolar epithelial reprogramming. * = *p* < 0.05. G) Expression of genes measured by qRT-PCR related to alveolar epithelial injury. H) Representative western blot of intracellular protein lysates from alveolospheres stimulated with CC+DMSO or FC+DMSO and treated with either nintedanib (FC+N10) or pirfenidone (FC+P10). Human lung tissue lysate as positive control. The signals were quantified as band intensity normalized to β-actin and expressed as percentages compared to the respective CC+DMSO sample which is set to 100 % expression. Data presented as means ± SD (FC+DMSO, FC+N10 and FC+P10). # = *p* < 0.05 by one sample t-test on the percentage intensity compared to a hypothetical value of 100 (CC+DMSO vs. FC+DMSO) and repeated measures one-way ANOVA test performed (FC+DMSO vs. FC+N10 or FC+P10). *n* = 3 batches of iAT2 cells. Blots are cropped from the same membrane and are obtained by sequential blotting as outlined in Supplemental Experimental Procedures. Brightness and contrast has been enhanced. For C, E, F and G: Expression of genes normalized to the average CT value of the reference genes *GAPDH, TBP* and *HPRT1.* Fold changes (2^-ΔΔCT) calculated by comparison to the average ΔCT value of the CC+DMSO population. Data presented as means ± SD. Significance tested by repeated measures one-way ANOVA, if significant with Dunnett’s multiple comparisons test (comparisons to FC+DMSO). *n* = 3 batches of iAT2 cells. See also Fig. S5, S6 and S7.

Intriguingly, treatment of the alveolospheres with 10 μM nintedanib partially prevented the induction of the dense morphology by the FC, which was not seen with pirfenidone (Fig. 6B). To further determine the molecular changes behind this morphological effect, we first evaluated if there were effects of the treatment on the FC-induced production of ECM and pro-fibrotic mediators (Fig. 6C-E). Treatment with 10 µM of nintedanib significantly decreased gene expression of *FN1* (Fig. 6C), and a similar trend was observed for the protein secretion in the medium (*p* of ANOVA = 0.099) (Fig. 6D). In contrast to the effects described in published studies using PCLS from bleomycin-injured mice (27), we could not observe any decrease of *COL1A1* gene expression in the alveolospheres after any of the treatments (Fig. S7B). However, we did observe significant effects of nintedanib and pirfenidone on the pro-fibrotic mediators *TGFB1* and *IGFBP3* (Fig. 6E) and a trend towards a decrease of *MMP7* expression (Fig. S7B).

Since the expression of markers associated with alveolar epithelial reprogramming and senescence were not significantly altered upon withdrawal of the FC, we investigated if the therapies could prevent the changes in these markers (Fig. 6F-H and S7B). Treatment with nintedanib significantly reduced the expression of the EMT markers *VIM* and *CDH2* (Fig. 6F). We also observed trends towards decreased expression of the senescence-associated markers *CDKN1A* and *CDKN2A* as well as of the reprogramming-associated markers *KRT8* and *KRT17* after treatment (Fig. S7B). Interestingly, none of the drugs prevented the loss of surfactant protein expression induced by the FC (Fig. 6G-H), indicating that the AT2 phenotype was not rescued by the treatment. Altogether, this data shows that the alveolospheres respond to anti-fibrotic therapy and have the potential to serve as a relevant model for drug screening.

## Discussion

Our aim was to create a model of alveolar epithelial injury in IPF adaptable to drug discovery. We have shown that IPF can be modeled *in vitro* by exposing iAT2 cells to a pro-fibrotic milieu mimicked by the FC. The virtue of iAT2 cells is that they can be derived from accessible adult somatic cells (17, 18, 50), which does not require organ donation and is of less ethical concern than the isolation of ESCs. The expandability of the iAT2 cells in the miniaturized culture format is a significant benefit as it enables our system to be used for drug screening and precise genome editing using CRISPR/Cas9 (51). We have used the same concentrations of the components of the FC as reported initially (26) which have shown effects on various cell types in PCLS from several human lung tissue donors (27). We have been able to study the effects of the FC on AT2-like cells which are isolated from non-epithelial cell types, a challenging aspect to study in systems like the *ex vivo* PCLS model or the *in vivo* bleomycin model. Another study has similarly explored the possibility of inducing fibrotic changes with an IPF-relevant cocktail (IPF-RC) in an iPSC-derived alveolar air-liquid interface model (22). However, the IPF-RC was different to the FC in composition and concentrations, and required a considerable longer stimulation of 15 days which is not desirable for rapid high throughput drug screening assays.

The shifts in the distribution and identities of lung epithelial cells in IPF is a relatively recent observation due largely to the availability of single-cell RNA-seq data from human IPF lungs (5, 8–10). Thus, models in which these aberrant cell identities can be studied are only beginning to emerge. Other studies have modeled fibrotic alveolar epithelial injury in ESC/iPSC-derived alveolar organoids (19, 21, 23, 24, 48), although these models have required co-cultures with non-epithelial cells and therefore limit the possibility to study the effects of defined factors on alveolar epithelial cells as is possible in our model. It is encouraging that the FC induces expression of known biomarkers in IPF such as *MMP7* (31) and *IGFBP3* (52), and a pro-fibrotic signature including the secretion of ECM and aberrant expression of mesenchymal markers. In particular, the observation that the FC induced an aberrant basaloid-like phenotype in our system will allow us to study the induction of this rare population by using defined stimulants without the need of co-cultures. The recent studies of epithelial reprogramming together with ours indicate that the aberrant epithelial subpopulations present in the IPF lung may be due to reprogramming of the distal epithelium as opposed to migration of proximal epithelial progenitors into the distal airspaces (53).

We also explored if our model can be used in drug screening by using a prophylactic treatment approach with the two approved therapies nintedanib and pirfenidone. Although prophylaxis in the clinic is a challenging concept as IPF is not detected at onset, this rationale could be appropriate for preventing progression of injury or after exacerbations. Interestingly, although we observed effects of nintedanib on markers associated with fibrosis and EMT in line with published studies (27, 54), we did not observe any rescue of the surfactant protein expression in contrast to other studies using primary AT2 cells from bleomycin-injured mice and human PCLS (27). The variation in responses could be due to the difference in species, culture on plastic as opposed to Matrigel, and effects of other cell types present in the PCLS. We have in this study given an overview of the responses to anti-fibrotic therapy as a proof of concept that our system has potential for use in drug screening, and our results highlight that there is a need for novel anti-fibrotic compounds which target the alveolar epithelium (55). Moreover, the fact that cryopreserved alveolospheres are fit for such experiments allows for performing drug screens on demand.

In conclusion, we have shown that stimulation of iAT2 cells with the FC is a system that models key features of human IPF. The model is suitable for studying fibrotic alveolar epithelial reprogramming and for use in drug discovery.

## Experimental procedures

### Cells and human lung tissue

The r-iPSC1J cell line was reprogrammed from newborn foreskin BJ fibroblasts (ATCC, CRL2522) acquired by AstraZeneca in compliance with the ATCC materials transfer agreement as previously described (56). The ethical consent form for the BJ fibroblast line was requested by AstraZeneca from ATCC but was not available. Therefore, the generation of the r-iPSC1J cell line was reviewed and supported by the AstraZeneca Human Biological Sample Governance Team.

The use of human lung tissue was approved by the local ethics committee in Lund, Sweden (Regionala Etikprövningsnämnden, Lund), Dnr 2013/253. All donors provided informed consent and the experiments were conducted in accordance with the Declaration of Helsinki.

### Fibrosis cocktail stimulation of alveolospheres

Thawed alveolospheres in passage 2-3 were resuspended and cultured in growth factor-reduced Matrigel (Corning, 354230) in 96-well plates with CK+DCI medium (Table S1 and S4) as described previously (17, 18) for 2-7 days before the first single cell passage. Single iAT2 cells in passages 4-6 (2-3 passages post thaw) were plated at a density of 100 cells/μl and formed organoids for 14 days. To induce fibrosis, alveolospheres were stimulated with the addition of the FC consisting of 5 ng/ml recombinant TGF-β (R&D Systems, 240-B-002/CF), 10 ng/ml PDGF-AB (GIBCO, PHG0134), 10 ng/ml TNF-α (R&D Systems, 410-MT-010/CF), and 5 μM LPA (Cayman Chemical, 62215) or the CC (diluents of components) to the CK+DCI medium for 72 h. For the FC withdrawal experiment, iAT2 cells were plated at a density of 50 cells/μl to avoid overgrowth due to longer culture time. For drug treatment, alveolospheres were stimulated with the FC with simultaneous addition of DMSO (control) and either nintedanib (Carbosynth, FM34144) or pirfenidone (Carbosynth, FM341441402) in concentrations 0.1 – 10 µM for 72 h.

### Statistics

The *n* numbers represent samples from differentiations performed at separate time points (batches) and data is presented as mean ± standard deviation (SD). The statistical tests are described separately in the figure legends. *p* values of < 0.05 were considered significant. Statistical tests and graphs were generated using GraphPad Prism v9 unless otherwise stated.

### Data and code availability

The RNA-seq data generated in this study is deposited in ArrayExpress with accession ID: E-MTAB-11676.

## Supporting information

List_S1

List_S2

List_S3

List_S4

List_S5

List_S6

List_S7

List_S8

List_S9

List_S10

List_S11

## Acknowledgements

We are grateful to Hani N. Alsafadi (Lund University) for his input on the immunofluorescence and deconvolution and to Johan Mattsson and Ulf Gehrmann (AstraZeneca) for performing the human lung tissue lysis. We thank Source Bioscience for the library preparation and sequencing services. Finally, we thank the donors of the lung tissue and the Sahlgrenska University Hospital for the technical support. The Knut and Alice Wallenberg foundation, the Medical Faculty at Lund University and Region Skåne are acknowledged for generous financial support (D.E.W.).

## Author contributions

Conceptualization, V.P., D.E.W. and L.A.M.; Methodology, V.P., K.Ö., M.T. and C.O-S.; Validation, V.P., K.Ö.; Formal analysis, V.P., S.J.M., K.Ö., M.T.; Investigation, V.P., K.Ö., M.T. and C.O-S.; Writing-Original Draft, V.P., P.H., D.E.W. and L.A.M.; Writing-Review & Editing, V.P., K.Ö., M.T., S.J.M., C.O-S., P.H., D.E.W. and L.A.M; Supervision: P.H., D.E.W. and L.A.M., Funding acquisition, D.E.W. and L.A.M.

---

**Figure S1:**
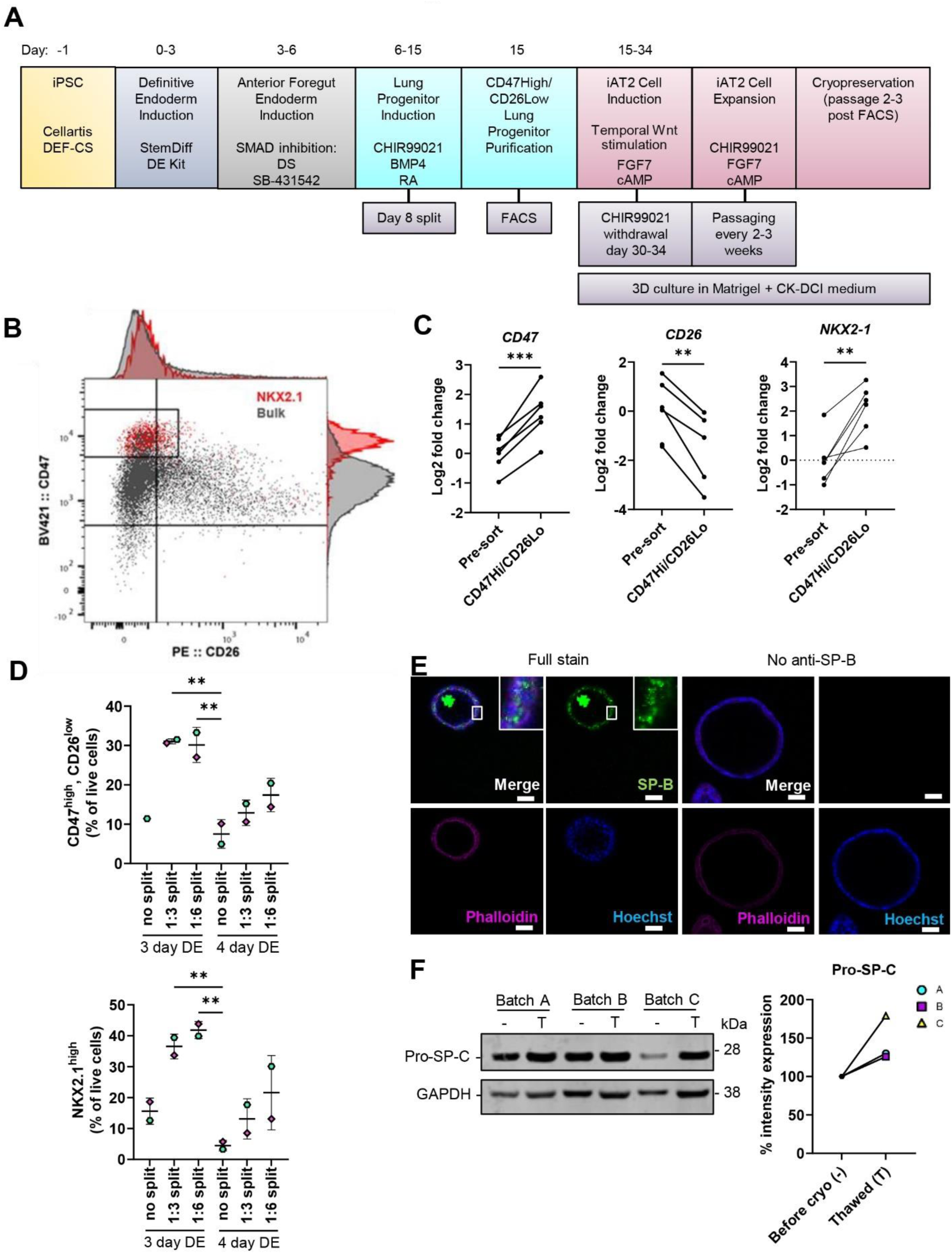
Derivation and validation of iAT2 cells. Related to Fig. 1. A) Differentiation scheme of iPSCs to iAT2 cells (1, 2). B) Representative flow cytometric analysis of the gating strategy for sorting the CD47^high^/CD26^low^ population, in which NKX2.1 protein expression was enriched at day 15 of differentiation. Red = NKX2.1^+^ cells, grey = bulk cell population (all single live cells). C) Gene expression validating the *NKX2-1* enrichment in the sorted CD47^high^/CD26^low^ populations measured by qRT-PCR. The expression of the gene in each sample was normalized to the average CT value of the reference genes *18S*, *TBP* and *HPRT1*. Fold changes (2^-ΔΔCT) were calculated by comparison to the average ΔCT value of the bulk lung progenitor population (pre-sort). *** = *p* < 0.001, ** = *p* < 0.01 by paired, two-tailed t-test. *n* = 6 batches of differentiated cells. D) Percentage of CD47^high^/CD26^low^ cells and NKX2.1^high^ cells by flow cytometry after different timings of DE induction and splits at the lung progenitor stage. Data presented as means ± SD. ** = *p* < 0.01 by ordinary one-way ANOVA with Dunnett’s multiple comparisons test (comparisons to 4 day DE, no split). Unpaired, two-tailed t-test performed (3 day DE 1:3 split vs. 3 day DE 1:6 split). *n* = 2 batches of differentiated cells. E) Immunofluorescence of pro- and mature form of SP-B in alveolospheres. No anti-SP-B as negative control for the staining. Scale bar = 50 µm. Brightness and contrast has been enhanced. F) Western blot of pro-SP-C expression in 3 batches of differentiated alveolospheres, before cryopreservation at passages 3-5 (-) and at 2 passages after thawing (T). Intensity expression (%) of pro-SP-C in alveolospheres before cryopreservation (set to 100 %) and after cryopreservation, normalized to the intensity of GAPDH. Blots are cropped from the same membrane. Brightness and contrast has been enhanced.

**Figure S2:**
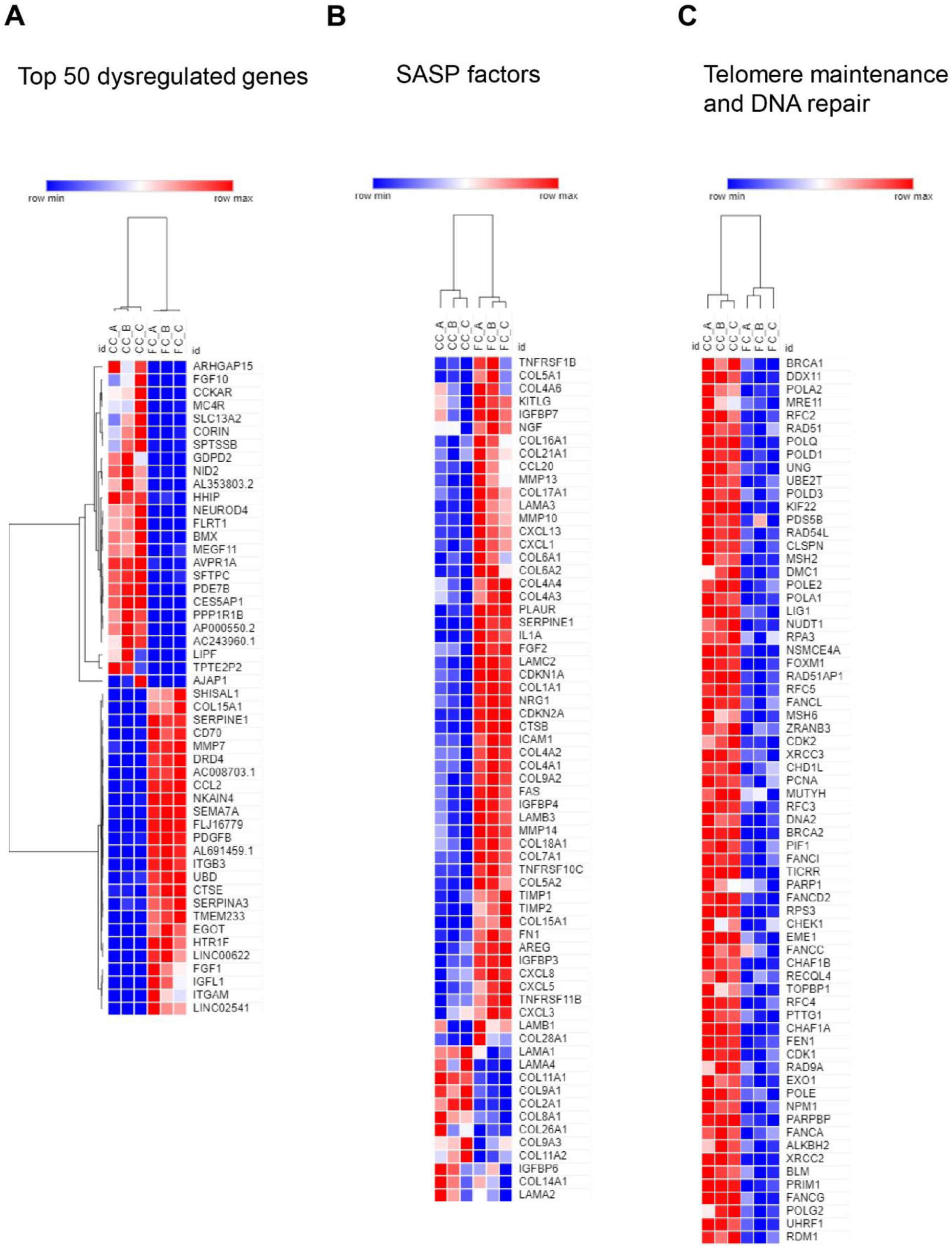
Dysregulated transcripts in alveolospheres stimulated with the fibrosis cocktail. Related to Fig. 1. A) Heatmap representation of top 50 dysregulated genes in FC-stimulated alveolospheres compared to CC stimulation by absolute fold change. *p(adj) <* 0.05, absolute log2 fold change ≥ 0.7. B) Heatmap representation of selected significantly dysregulated SASP factors in FC-stimulated alveolospheres compared to CC stimulation by absolute fold change. *p(adj) <* 0.05, absolute log2 fold change ≥ 0.7. C) Heatmap representation of selected significantly dysregulated genes related to GO_BP:0000722: telomere maintenance via recombination and GO:0006281: DNA repair in FC-stimulated alveolospheres compared to CC stimulation by absolute fold change. *p(adj) <* 0.05, absolute log2 fold change ≥ 0.7.

**Figure S3:**
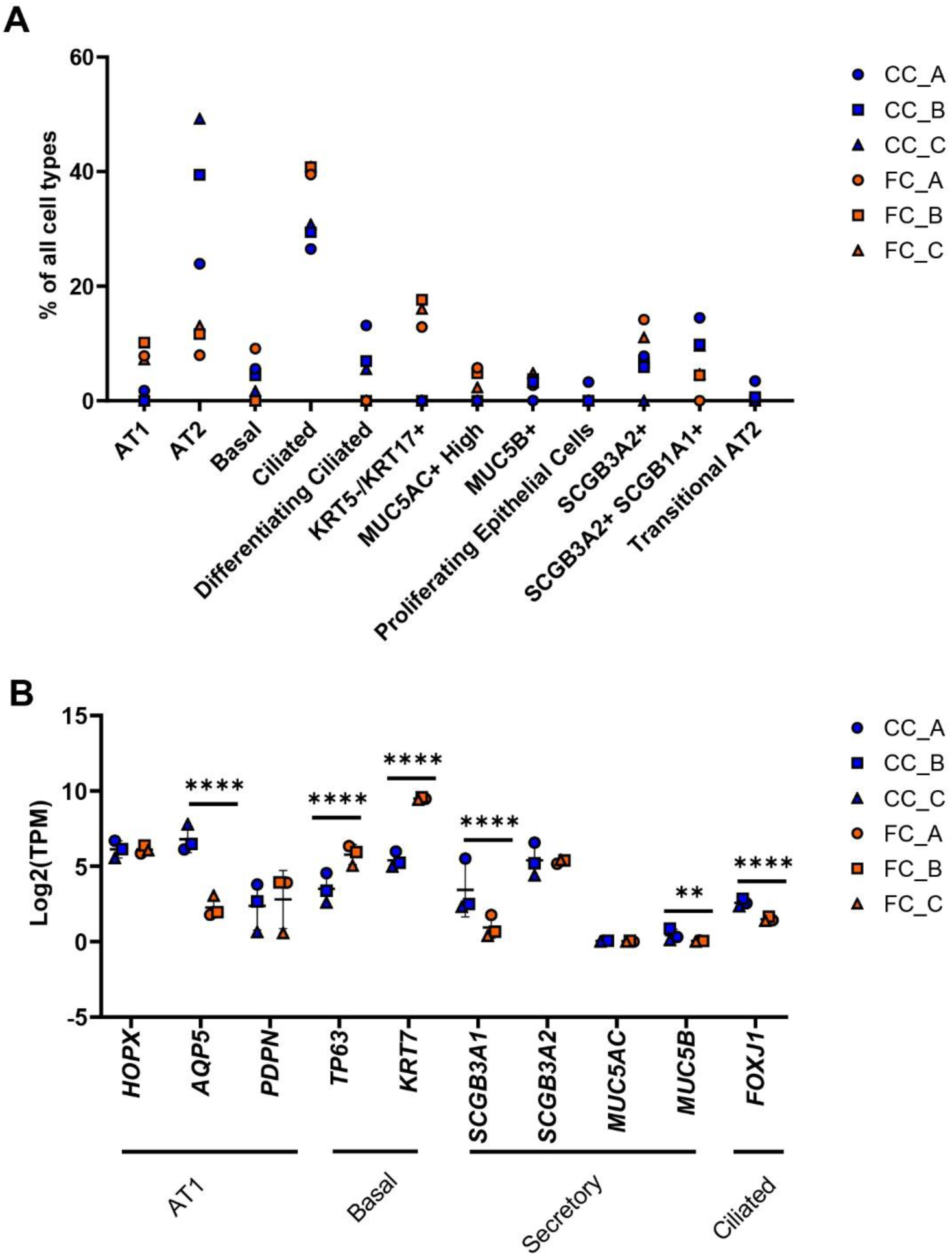
Deconvolution of RNA-seq data from alveolospheres stimulated with the fibrosis cocktail. Related to Fig. 3. A) All epithelial cell proportion estimates by Bisque in CC-stimulated or FC-stimulated alveolospheres. *n =* 3 batches of iAT2 cells. B) Expression of selected epithelial genes specific for AT1, basal, secretory and ciliated cells. Data presented as means ± SD. **** = *p(adj) <* 0.0001, ** = *p(adj) <* 0.01. *n =* 3 batches of iAT2 cells.

**Figure S4:**
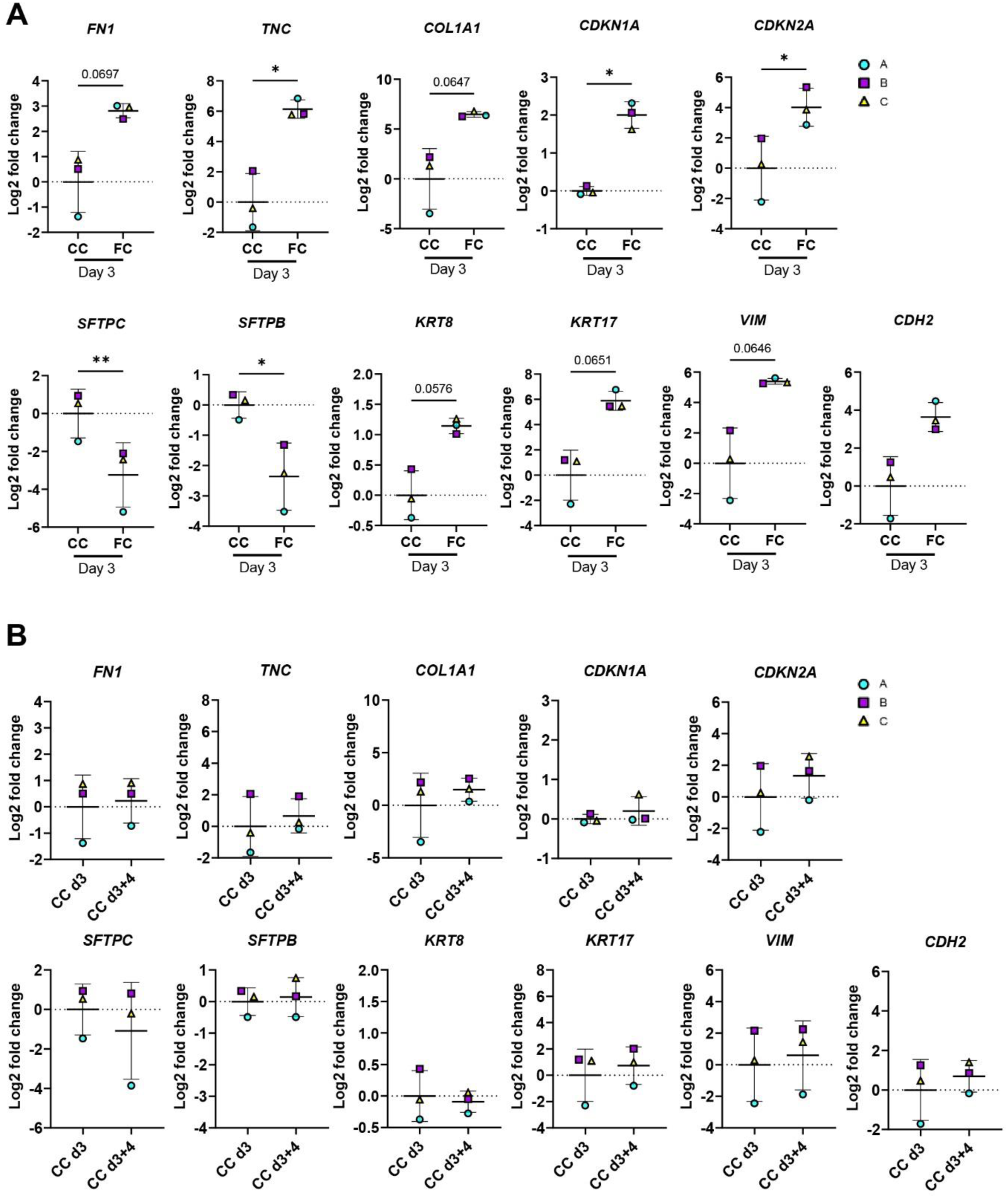
Assessment of markers for evaluation of fibrosis cocktail effects after withdrawal. Related to Fig. 5. A) Expression of genes measured by qRT-PCR. The expression of the gene in each sample was normalized to the average CT value of the reference genes *GAPDH, TBP* and *HPRT1.* Fold changes (2^- ΔΔCT) were calculated by comparison to the average ΔCT value of the CC day 3 population. Data presented as means ± SD. ** = *p* < 0.01, * = *p* < 0.05 by paired, two-tailed t-test. *n =* 3 batches of iAT2 cells. B) Baseline expression of genes in the CC samples at day 3 and after withdrawal at day 3+4, measured by qRT-PCR. The expression of the gene in each sample was normalized to the average CT value of the reference genes *GAPDH, TBP* and *HPRT1.* Fold changes (2^-ΔΔCT) were calculated by comparison to the average ΔCT value of the CC day 3 population. Data presented as means ± SD. Paired, two-tailed t-test performed. *n =* 3 batches of iAT2 cells.

**Figure S5:**
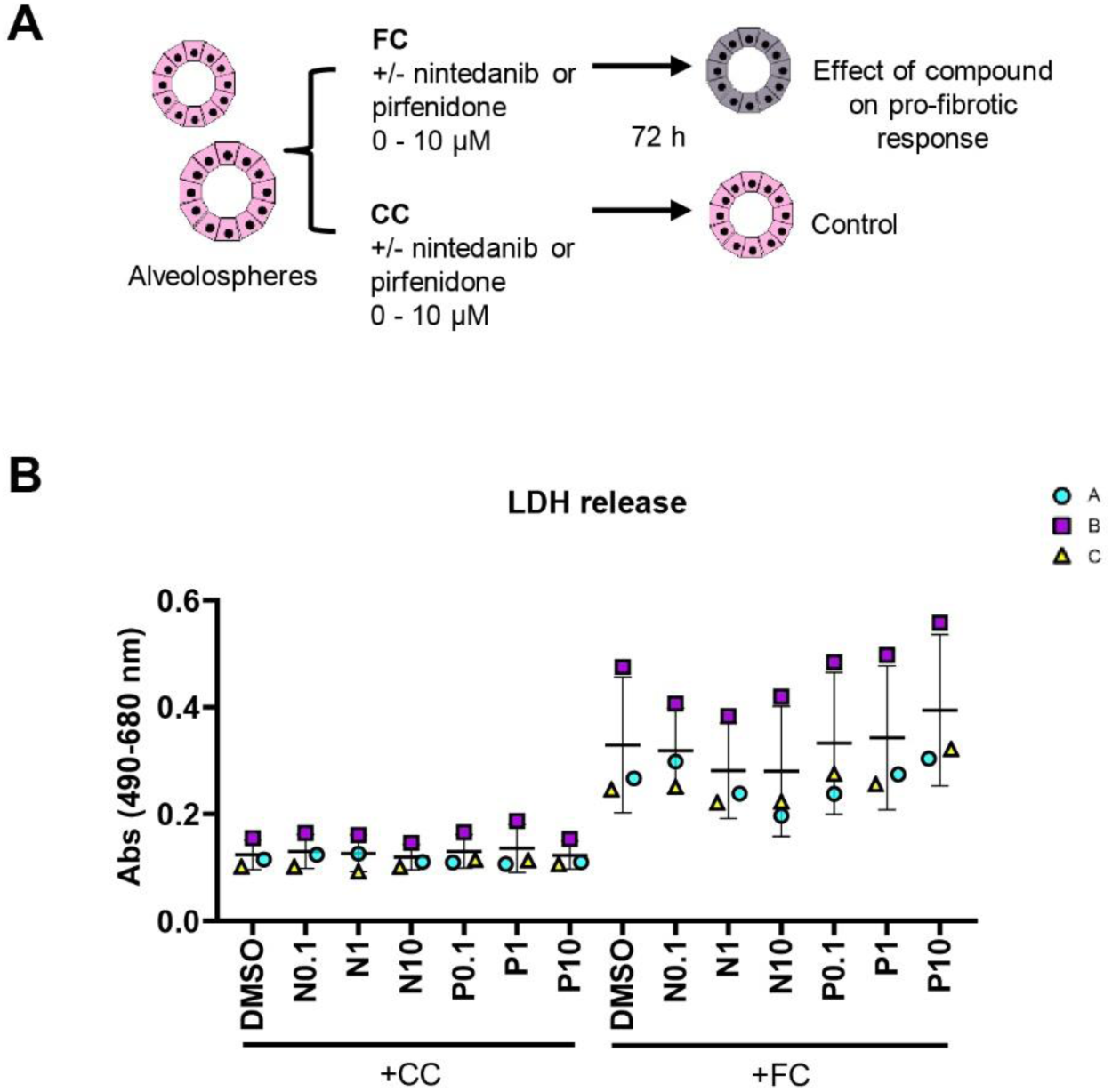
Cytotoxicity following treatment with anti-fibrotic compounds of alveolospheres. Related to Fig. 6. A) Schematic illustrating prophylactic treatment of alveolospheres stimulated with either the CC or the FC in the presence or absence of nintedanib or pirfenidone. B) LDH release detected in secreted medium of alveolospheres stimulated with either the CC or the FC in the presence or absence of nintedanib (N) or pirfenidone (P). Data presented as means of absorbance values of eight technical replicates for each batch ± SD. Repeated measures one-way ANOVA (comparisons of CC+N or P vs. CC+DMSO, and FC+N or P vs. FC-DMSO) performed. *n =* 3 batches of iAT2 cells.

**Figure S6:**
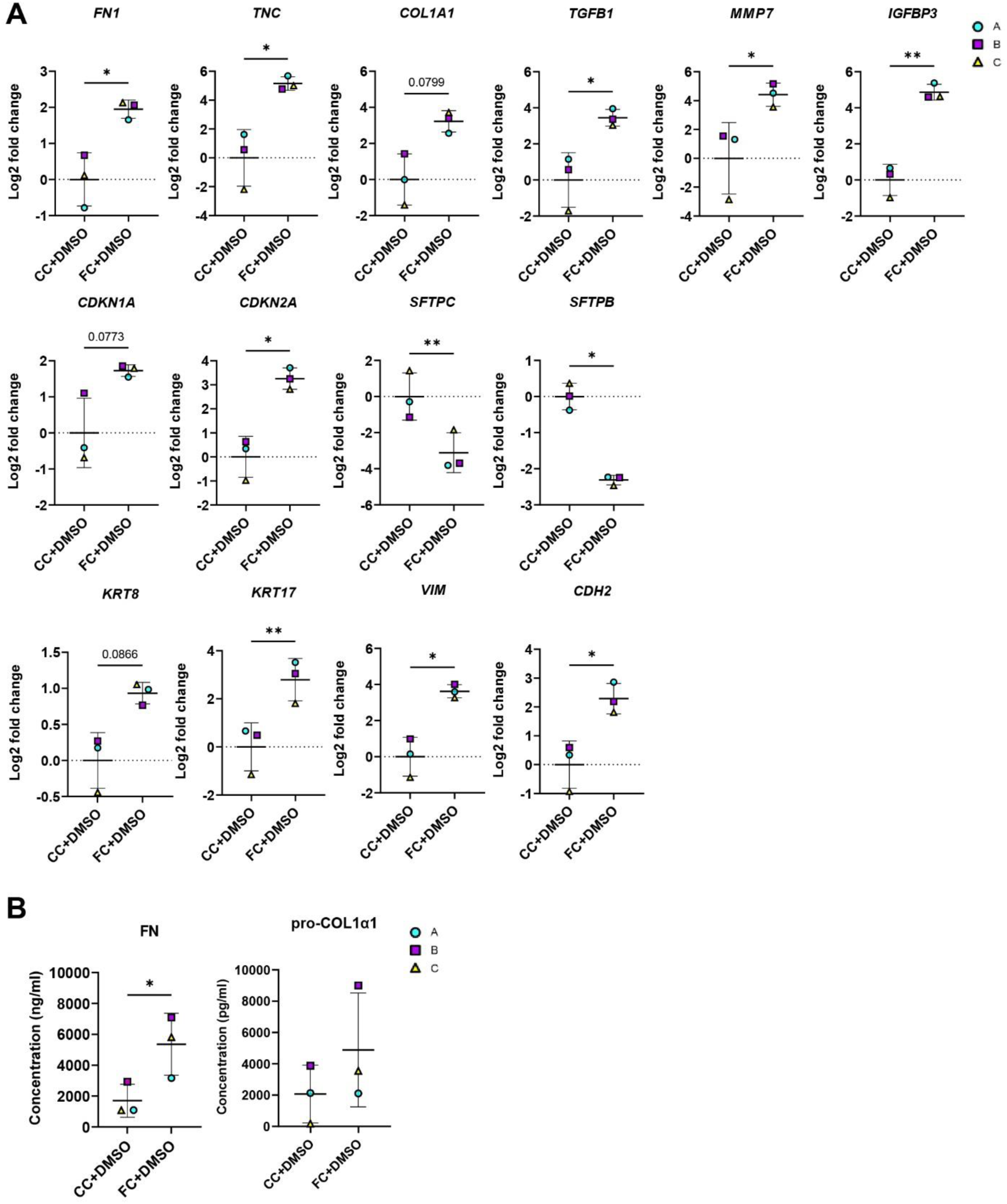
Effects of FC or CC with DMSO on markers for evaluation of anti-fibrotic treatment. Related to Fig. 6. A) Expression of genes measured by qRT-PCR. The expression of the gene in each sample was normalized to the average CT value of the reference genes *GAPDH, TBP* and *HPRT1.* Fold changes (2^- ΔΔCT) were calculated by comparison to the average ΔCT value of the CC+DMSO population. Data presented as means ± SD. ** = *p* < 0.01, * = *p* < 0.05 by paired, two-tailed t-test. *n* = 3 batches of iAT2 cells. B) Secreted protein concentrations of fibronectin and pro-collagen 1α1 in the medium measured by ELISA. Data presented as means ± SD. * = *p* < 0.05 by paired, two-tailed t-test. *n* = 3 batches of iAT2 cells. Note: Values of FC+DMSO are the same as shown in Fig. 6 to allow for comparisons.

**Figure S7:**
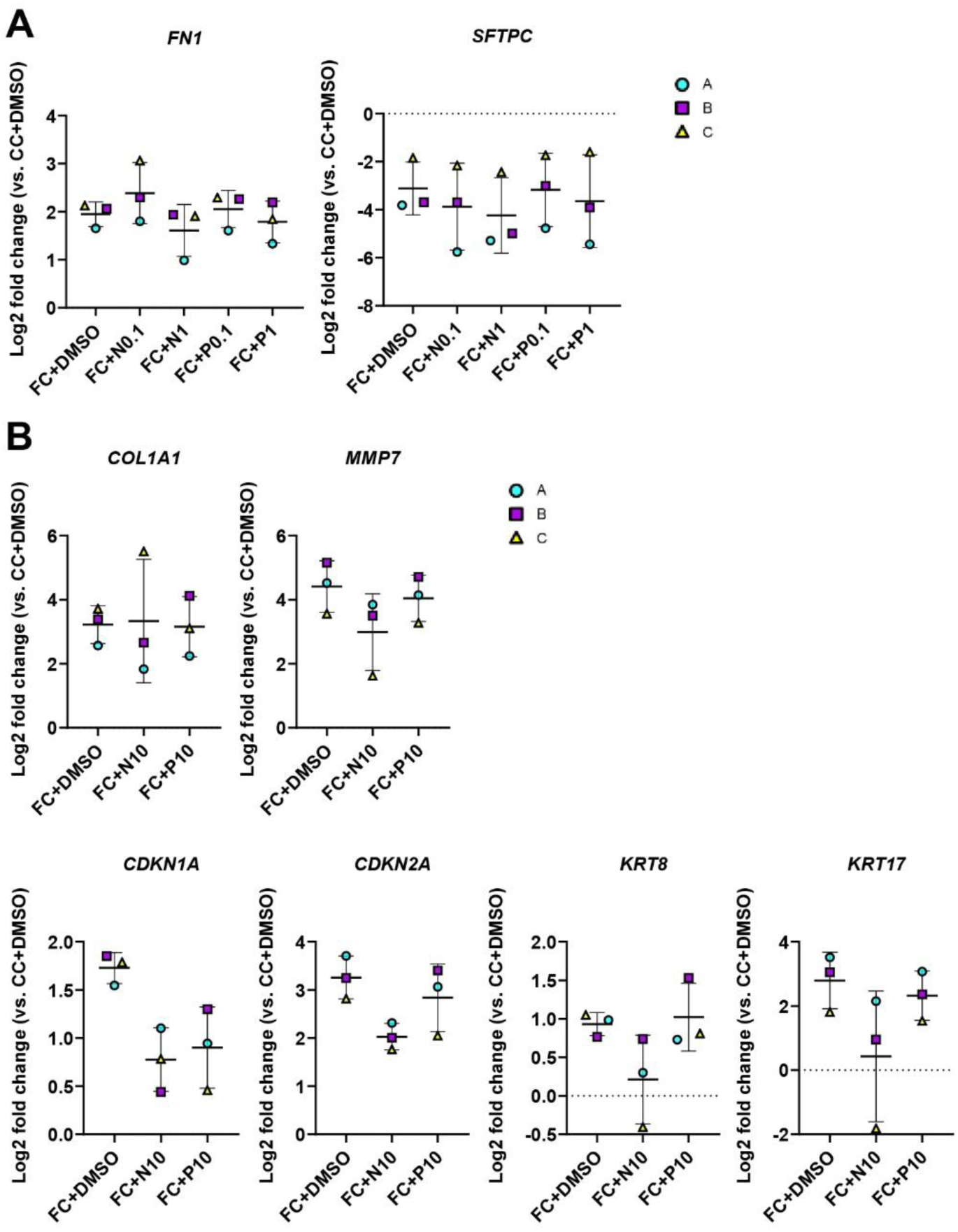
Effects of anti-fibrotic compounds on markers dysregulated by the fibrosis cocktail. Related to Fig. 6. A) Expression of genes measured by qRT-PCR. The expression of the gene in each sample was normalized to the average CT value of the reference genes *GAPDH, TBP* and *HPRT1.* Fold changes (2^- ΔΔCT) were calculated by comparison to the average ΔCT value of the CC+DMSO population. Data presented as means ± SD. Repeated measures one-way ANOVA (comparisons to FC+DMSO) performed. *n* = 3 batches of iAT2 cells. B) Expression of genes altered by the FC where treatment effects were non-significant, measured by qRT-PCR. Expression of genes normalized to the average CT value of the reference genes *GAPDH*, *TBP* and *HPRT1*. Fold changes (2^-ΔΔCT) calculated by comparison to the average ΔCT value of the CC+DMSO population. Data presented as means ± SD. Repeated measures one-way ANOVA (comparisons to FC+DMSO) performed. *n* = 3 batches of iAT2 cells. Note: Values of FC+DMSO are the same as shown in Fig. 6 and Fig. S6 to allow for comparisons.

## Supplemental Lists

List S1: Differentially expressed genes in alveolospheres + FC vs. CC

List S2: Upregulated GO:Biological process

List S3: Downregulated GO: Biological process

List S4: Upregulated GO: Cellular component

List S5: Downregulated GO: Cellular component

List S6: Upregulated GO: Molecular function

List S7: Downregulated GO: Molecular function

List S8: Upregulated KEGG Pathway

List S9: Downregulated KEGG Pathway

List S10: Common upregulated Venn

List S11: Common downregulated Venn

---

**Table S1:** Medium components for cSFDM medium.

| Component | Final concentration | CAT# | Vendor |
| --- | --- | --- | --- |
| **Iscove's Modified Dulbecco's Medium (IMDM)** | 75 % | 12440053 | Thermo Fisher |
| **Ham's F-12 Nutrient Mix** | 25 % | 11765054 | Thermo Fisher |
| **Bovine Albumin Fraction V (7,5%)** | 0.05 % | 15260037 | Thermo Fisher |
| **GlutaMAX 100X** | 1X | 35050061 | Thermo Fisher |
| **N-2 supplement 100X** | 0.5X | 17502048 | Thermo Fisher |
| **B-27 supplement 50X** | 0.5X | 17504044 | Thermo Fisher |

**Table S2:** Medium components for DS/SB medium, added to cSFDM.

| Component | Final concentration | CAT# | Vendor |
| --- | --- | --- | --- |
| **L-ascorbic acid** | 50 µg/ml | A4544 | Sigma-Aldrich |
| **1-thioglycerol** | 0.45 mM | M6145 | Sigma-Aldrich |
| **Dorsomorphin (DS)** | 2 µM | 3093 | Tocris |
| **SB-431542** | 10 µM | 1614 | Tocris |
| **Y-27632**<br>**(only when passaging)** | 10 µM | 688000 | Calbiochem |

**Table S3:** Medium components for CBRa medium, added to cSFDM.

| Component | Final concentration | CAT# | Vendor |
| --- | --- | --- | --- |
| **L-ascorbic acid** | 50 µg/ml | A4544 | Sigma-Aldrich |
| **1-thioglycerol** | 0.45 mM | M6145 | Sigma-Aldrich |
| **CHIR99021** | 3 µM | 4423 | Tocris |
| **Bone morphogenetic protein 4 (BMP4), human recombinant** | 10 ng/ml | 314-BP | R&D Systems |
| **Retinoic acid (RA)** | 100 nM | 0695 | Tocris |
| **Y-27632**<br>**(only when passaging)** | 10 µM | 688000 | Calbiochem |

**Table S4:** Medium components for CK+DCI medium, added to cSFDM.

| Component | Final concentration | CAT# | Vendor |
| --- | --- | --- | --- |
| **L-ascorbic acid** | 50 µg/ml | A4544 | Sigma-Aldrich |
| **1-thioglycerol** | 0.45 mM | M6145 | Sigma-Aldrich |
| **CHIR99021** | 3 µM | 4423 | Tocris |
| **Fibroblast growth factor 7 (FGF7), human recombinant** | 10 ng/ml | 251-KG | R&D Systems |
| **Dexamethasone** | 50 nM | D4902 | Sigma-Aldrich |
| **3-Isobutyl-1-methylxanthine (IBMX)** | 0.1 mM | I5879 | Sigma-Aldrich |
| **8-Bromoadenosine 3',5'-cyclic monophosphate (cAMP)** | 0.1 mM | B5386 | Sigma-Aldrich |
| **Y-27632**<br>**(only when passaging)** | 10 µM | 688000 | Calbiochem |
| **Primocin**<br>**(only during first week after FACS)** | 200 ng/ml | NC9141851 | Invivogen |

**Table S5:** Primary antibodies used for FACS, flow cytometry, western blotting and immunofluorescence.

| Target | Species | CAT# | Vendor | Dilution | Application |
| --- | --- | --- | --- | --- | --- |
| CD26-PE conj. | Mouse | 302705 | Biolegend | 0.025 µg/<br>5 x 10<sup>6</sup> cells | FACS |
| CD47-BV421 conj. | Mouse | 323115 | Biolegend | 0.05 µg/<br>5 x 10<sup>6</sup> cells | FACS |
| IgG2aκ-PE conj.<br>isotype control | Mouse | 400211 | Biolegend | 0.025 µg/<br>5 x 10<sup>6</sup> cells | FACS |
| IgG1κ-BV421 conj.<br>isotype control | Mouse | 562438 | BD Biosciences | 0.05 µg/<br>5 x 10<sup>6</sup> cells | FACS |
| NK2 Homeobox 1<br>(NKX2.1) | Mouse | ab72876 | Abcam | 3.2 µg /<br>2 x 10<sup>5</sup> cells | Flow cytometry |
| Pro-surfactant<br>protein C (SP-C) | Rabbit | AB3786 | Merck | 1:500 | Western blotting |
| Pro + mature<br>surfactant protein<br>B (SP-B) | Rabbit | ab40876 | Abcam | 1:1000 | Western blotting |
| β-Actin | Goat | ab8229 | Abcam | 1:1000 | Western blotting |
| Glyceraldehyde-3-phosphate<br>dehydrogenase<br>(GAPDH) | Mouse | ab9482 | Abcam | 1:1000 | Western blotting |
| Pro-SP-C | Rabbit | AB3786 | Merck | 1:500 | Immunofluorescence |
| Pro + mature SP-B | Rabbit | ab40876 | Abcam | 1:500 | Immunofluorescence |
| E-Cadherin (ECAD) | Mouse | 610181 | BD Biosciences | 1:100 | Immunofluorescence |
| Collagen 1 (COL1) | Rabbit | ab34710 | Abcam | 1:100 | Immunofluorescence |
| Vimentin (VIM) | Rabbit | 5741 | Cell Signaling | 1:100 | Immunofluorescence |
| Keratin 8 (KRT8) | Mouse | UM500001 | Thermo Fisher | 1:50 | Immunofluorescence |
| Keratin 17 (KRT17) | Rabbit | HPA000452 | Merck | 1:100 | Immunofluorescence |

**Table S6:** TaqMan® assays used for qRT-PCR.

| Target | Assay ID | Vendor |
| --- | --- | --- |
| CD47 | Hs00179953_m1 | Applied Biosystems |
| CD26 | Hs00897386_m1 | Applied Biosystems |
| NKX2-1 | Hs00968940_m1 | Applied Biosystems |
| SFTPB | Hs01090667_m1 | Applied Biosystems |
| SFTPC | Hs00161628_m1 | Applied Biosystems |
| KRT8 | Hs01595539_g1 | Applied Biosystems |
| KRT17 | Hs00356958_m1 | Applied Biosystems |
| FN1 | Hs01549976_m1 | Applied Biosystems |
| TNC | Hs01115665_m1 | Applied Biosystems |
| COL1A1 | Hs00164004_m1 | Applied Biosystems |
| CDH2 | Hs00983056_m1 | Applied Biosystems |
| VIM | Hs00958111_m1 | Applied Biosystems |
| CDKN1A | Hs00355782_m1 | Applied Biosystems |
| CDKN2A | Hs02902543_mH | Applied Biosystems |
| TGFB1 | Hs00998133_m1 | Applied Biosystems |
| IGFBP3 | Hs00181211_m1 | Applied Biosystems |
| MMP7 | Hs01042796_m1 | Applied Biosystems |
| TBP | Hs00427620_m1 | Applied Biosystems |
| HPRT1 | Hs02800695_m1 | Applied Biosystems |
| GAPDH | Hs99999905_m1 | Applied Biosystems |
| 18S | Hs99999901_s1 | Applied Biosystems |

## Supplemental Experimental Procedures

### Human iPSC maintenance

iPSCs from the previously described line r-iPSC1J (3) were cultured using the Cellartis® DEF-CS™ 500 Culture System (Clontech, Y30010) according to manufacturer’s instructions.

### Differentiation of human iPSCs to iAT2 cells

Directed differentiation of iPSCs to iAT2 cells was performed as previously described (1, 2, 4) with minor modifications; briefly, iPSCs were seeded at 6 million cells per Matrigel-coated (Corning, 354277) T25 flask in Cellartis® DEF-CS™medium (Clontech, Y30010) supplemented with 10 µM Y-27632 (Calbiochem, 688000). 24 hours later, the cells were differentiated to definitive endoderm (DE) using the commercial STEMdiff™ Definitive Endoderm Kit (Stemcell Technologies, 05110) for 3 days. For the next differentiation steps, cells were cultured in chemically defined media in complete serum-free differentiation medium (cSFDM) (Table S1). Following the DE induction, medium was switched to “DS/SB” medium for anterior foregut endoderm induction (Table S2) for another 3 days. At day 6 of differentiation, the medium was switched to “CBRa” medium (Table S3) until day 15 of differentiation. At day 8 of differentiation, a 1:3 ratio cell split was introduced. For iAT2 cell differentiation and maturation, sorted CD47^high^/CD26^low^ lung progenitor cells according to the previously published protocols (1, 2, 4) were cultured in 20 µl drops of growth factor-reduced Matrigel (Corning, 354230) in 96-well plates with lung epithelium maturation medium “CK+DCI” (Table S4) and passaged by single cell dissociation every 2-3 weeks as described previously using dispase (Corning, 354235) (1, 2). At passage 2-3, alveolospheres were cryopreserved in freezing medium consisting of 90 % embryonic stem cell (ESC)-qualified fetal bovine serum (FBS) (Thermo Fisher, 16141061) and 10 % dimethyl sulfoxide (DMSO) (Sigma-Aldrich, D8418) according to the previously published protocols (1, 2).

### Fluorescence-activated cell sorting (FACS)

At day 15 of differentiation, lung progenitor cells were dissociated, resuspended in FACS buffer containing 1x HBSS (-/-) (Gibco, 14175-053), 2% ESC-qualified FBS, 10 μM Y-27632 (Calbiochem, 688000) and 200 ng/ml Primocin (Invivogen, NC9141851) and stained for FACS analysis as earlier described (4) using primary antibodies or isotype controls as outlined in Table S5. Cell sorting was performed on a Sony SH800 Cell Sorter (Sony Biotechnology) and analysis acquired using the accompanying software (Sony Biotechnology). For intracellular staining of NKX2.1, FACS buffer containing phosphate-buffered saline (PBS) (-/-) (Thermo Fisher, 10010-015), 2% FBS (Thermo Fisher, 10270-106) and 2 mM ethylenediaminetetraacetic acid disodium salt dihydrate (EDTA) (Sigma Aldrich, E4884) was used and prior to antibody staining, cells were first fixed with Fix Buffer I (BD Biosciences, 557870) and permeabilized with Perm Buffer III (BD Biosciences, 558050). Analysis and plots were generated using the software FlowJo v.10 (BD Biosciences).

### Reverse transcriptase quantitative PCR (qRT-PCR)

RNA from alveolospheres was isolated using TRIzol™ Reagent (Invitrogen, 15596026) according to manufacturer’s instructions. mRNA was further purified from the aqueous phase using the RNeasy Micro Kit (Qiagen, 74004) according to manufacturer’s instructions. Reverse transcription was performed using the High-Capacity cDNA Reverse Transcription Kit (Applied Biosystems, 4368813) according to manufacturer’s instructions on a Peltier Thermal Cycler PTC-200 (MJ Research). For qRT-PCR analysis, individual TaqMan® Gene Expression Assays (Applied Biosystems) were used with an amount of 1.3-2 ng cDNA per assay and TaqMan® Low Density Arrays (Applied Biosystems, 4342253) were used with a total amount of 200 ng cDNA per sample (2 ng cDNA per assay) and run for 45 cycles using the QuantStudio7 system (Applied Biosystems). CT values above 40 were considered as no expression. Gene expression was normalized to the appropriate endogenous controls as indicated and expressed as fold changes calculated using the 2^(−ΔΔCT) method relative to the corresponding control samples as denoted for each case. A list of all TaqMan® Gene Expression Assays used in this study is provided in Table S6.

### RNA-sequencing

RNA from alveolospheres was isolated as already described. Concentration and quality of the RNA was determined using the RNA 6000 Nano Assay on the Agilent Bioanalyzer (Agilent Technologies). Libraries were prepared by Source Bioscience (Nottingham, United Kingdom) using the NEBNext® Ultra™ II Directional RNA Library Prep Kit for Illumina® with mRNA module (New England Biolabs Inc., E7760) and sequenced (50 ng, 150bp paired-end) on a NovaSeq 6000 (Illumina).

### Bioinformatic analysis of RNA-sequencing data

The fastq files were processed with the bcbio-nextgen v 1.2.3 pipeline using Hisat2 (v2.2.0) as the aligner and Salmon (v1.1.0) as the pseudo-aligner. The reference genome was hg38, ensembl annotation release 99. In downstream analyses the TPM values generated from Salmon were used as the expression values. The differential expression was performed using the Salmon generated pseudo-counts processed with tximport in DESeq2 with apeglm shrinkage to generate log2 fold changes and multiple testing adjusted *p* values (*p(adj)*). Principle component analysis (PCA) was performed based on the top 1000 transcripts by coefficient of variation. To generate the list of differentially expressed genes (DEG), dysregulated transcripts were defined by an absolute log2 fold change of 0.7 and a FDR-adjusted p-value (*p(adj)) <* 0.05 (List S1). The gene ontology analysis was generated by The Database for Annotation, Visualization and Integrated Discovery (DAVID, v6.8, https://david.ncifcrf.gov/, accessed on 27 April 2021) based on the dysregulated transcripts defined by an absolute log2 fold change of 0.7 and *p(adj) <* 0.05 (Lists S2-S9), selected hits defined by *p(adj) <* 0.05 for graphs. The differentially expressed genes in alveolospheres stimulated with the FC were compared to publicly available data from human AT2 cells derived from IPF patients (5), GEO Accession: GSE94555. The gene sets were split into upregulated transcripts and downregulated transcripts and respective intersections were calculated. Venn-diagrams were produced by utilizing the following online tool: http://bioinformatics.psb.ugent.be/webtools/Venn/ and the complete lists of commonly upregulated and downregulated transcripts are presented in Lists S10 and S11, respectively. Heatmaps were generated using Morpheus, (https://software.broadinstitute.org/morpheus), linkage method average and one minus Pearson correlation. For heatmaps and expression plots log2(TPM) were used unless otherwise stated.

### Deconvolution analysis

Deconvolution of RNA-seq data was performed using the Bisque package in R v3.5.2 using the single-cell reference-based approach as already described (6, 7) based on a publicly available single-cell RNA-seq dataset (8), GEO Accession: GSE135893. Cells were filtered to only include epithelial cell populations from non-fibrotic donors and IPF patients.

### Western blot analysis

For intracellular protein retrieval, alveolospheres were dissociated from the Matrigel as described previously (1, 2) and lysed in RIPA Lysis and Extraction Buffer (Thermo Fisher, 89900) supplemented with cOmplete™ Protease Inhibitor Cocktail (Roche, 11697498001) and PhosSTOP™ inhibitors (Roche, 4906837001) on ice for 30 min. The lysate was then obtained by centrifugation at 15 000 x g for 15 min at 4 °C. Human lung tissue was homogenised for 2x 30 seconds at 25 Hz in ice cold PBS with cOmplete™ Protease Inhibitor Cocktail and PhosSTOP™ inhibitors, then lysed by addition of RIPA Lysis and Extraction buffer. The lysate was then obtained by centrifugation at 14 000 x g for 10 min at 4 °C. The total protein concentration was measured with the Pierce™ BCA Protein Assay Kit (Thermo Fisher, 23225). A total of 10-20 µg of intracellular protein was resolved on precast NuPAGE™ Novex™ 12 % Bis-Tris protein gels (Invitrogen, NP0343 and NP0341) and transferred to Invitrolon™ PVDF membranes (Invitrogen, LC2005). Blots were incubated with primary antibodies overnight at 4 °C as specified in Table S5. Species-specific secondary antibodies conjugated to infrared (IR) dyes of either 680 nm or 800 nm wavelengths (LiCOR Biosciences) were used at a dilution of 1:10 000 and the blots were visualized using the Odyssey Imaging System (LiCOR Biosciences), automatic setting of channel intensities, 169 µm resolution and quality-highest. The blots were blotted sequentially with anti-pro-SP-C (Merck, AB3786) with either anti-GAPDH (Abcam, ab9482) or anti-β-actin (Abcam, ab8229) and visualized, then re-blocked with blocking buffer containing TBS + 0.1 % Tween-20 (Takara, T9142) + 5 % BSA (Sigma, A6003) and re-incubated with anti-SP-B (Abcam, ab40876) overnight and visualized with secondary antibodies as described. The signals were quantified using the software Image Studio v4.0 (LiCOR Biosciences) on the original blots. For figures, brightness and contrast for the whole blot was adjusted for each channel individually, and regions of interest were cropped from the original blots for figure display.

### Immunofluorescence microscopy

Alveolospheres were fixed with 4 % paraformaldehyde for 1 h on ice and phase contrast images of fixed cultures were taken using the 4x objective with automatic focus on the Incucyte S3 (Sartorius). Alveolospheres were then permeabilized in blocking buffer containing 5 % normal goat serum (Vector Laboratories, S-1000) with 0.5 % Triton X-100 (Sigma) overnight at 4°C. The primary antibodies were diluted in blocking buffer and incubated at 4 °C overnight as specified in Table S5. Alveolospheres were then washed in blocking buffer and counter-stained with Hoechst33342 at 3.3 μg/ml (ThermoFisher, H3570) and the following secondary antibodies: goat anti-rabbit IgG Alexa Fluor 488 (Thermo Fisher, A11034, dilution 1:500), goat anti-mouse Alexa Fluor 594 (Thermo Fisher, A21125, dilution 1:500) and goat anti-mouse Alexa Fluor 647 (Thermo Fisher, A21241, dilution 1:500) in blocking buffer at 4 °C overnight. Visualization was performed by using a Zeiss LSM 880 laser scanning confocal microscope (Zeiss) at a resolution of 1024 px × 1024 px, 16-bit format using the Plan-Apochromat 20x/0.8 M27 objective and image processing was performed using the software ZEN v2.3 (Zeiss). The imaging settings were: for pro-SP-C/E-cadherin/Hoechst, laser strength: 633-10.0%, 488-1.0%, 405-1.0%, master gain: 800 (all lasers), digital gain: 633-3.0, 488-1.5, 405-1.0; for vimentin/E-cadherin/Hoechst, laser strength: 633-10.0%, 488-1.0%, 405-1.0%, master gain: 800 (all lasers), digital gain: 633-3.0; 488-10.0; 405-1.0; for keratin 17/keratin8/Hoechst, laser strength: 594-10.0%; 488-1.0%; 405-1.0%, master gain: 800 (all lasers), digital gain: 594-10.0; 488-2.5; 405-1.0; for collagen type 1/E-cadherin/Hoechst, laser strength: 633-10.0%; 488-1.0%; 405-1.0%, master gain: 800 (all lasers), digital gain: 633-3.0; 488-1.5; 405-1.0; for SP-B/Phalloidin/Hoechst, laser strength: 594-2.0%; 488-2.0%; 405-2.0%, master gain: 594-620; 488-569; 405-650, digital gain: 594-1.0; 488-1.0; 405-1.0. The images shown in the figures are maximum intensity projections of 10 vertical z-stack images (size 6.173 µm (6.175 µm for keratin 17/8)). For Fig S1E, single plane images were captured at 512 px × 512 px, 8-bit format using the LD Plan-Neofluar 20x/0.4 Korr M27 objective. For images in figures, brightness and contrast were adjusted for optimal signal display of each fluorophore and applied to the entire image, and were kept consistent within each staining combination and matched with no primary antibody controls. Display settings are: for pro-SP-C/E-cadherin/Hoechst, black: 0 (min 0), gamma: 1.00, white: 65535 (max 65535); for vimentin/E-cadherin/Hoechst, black: 20000 (min 0), gamma: 1.00, white: 65535 (max 65535), for keratin 17/keratin 8/Hoechst, black: 15000 (min 0), gamma: 1.00, white: 65535 (max 65535); for collagen type 1/E-cadherin/Hoechst, black: 25000 (min 0), gamma: 1.00, white: 65535 (max 65535); for SP-B/Phalloidin/Hoechst, black: 0, gamma: 1.00, white: 200 (max 255).

### Enzyme-linked immunosorbent assay (ELISA)

Secreted protein levels were assessed in conditioned medium from alveolospheres by ELISA for human fibronectin (Thermo Fisher, BMS2028), human tenascin (Abcam, ab213831) and human pro-collagen 1α1 (R&D Systems, DY6220-05) according to manufacturer’s instructions.

### LDH analysis

LDH analysis was performed using the CyQUANT™ LDH Cytotoxicity Assay Kit (Invitrogen, C20301) using fresh cell culture medium according to manufacturer’s instructions.

